# Architecture of the Mto1/2 microtubule nucleation complex

**DOI:** 10.1101/754457

**Authors:** Harish C. Thakur, Eric M. Lynch, Weronika E. Borek, Xun X. Bao, Sanju Ashraf, Juan Zou, Juri Rappsilber, Atlanta G. Cook, Kenneth E. Sawin

## Abstract

Proteins that contain a Centrosomin Motif 1 (CM1) domain are key regulators of *γ*-tubulin complex-dependent microtubule nucleation, but how they are organized in higher-order structures is largely unknown. Mto1[bonsai], a truncated functional version of the *Schizosaccharomyces pombe* CM1 protein Mto1, interacts with Mto2 to form an Mto1/2[bonsai] complex *in vivo*. Here we show that recombinant Mto1/2[bonsai] forms higher-order multimers *in vitro* and that Mto2 alone can also multimerize. We demonstrate that Mto2 multimerization involves two separate homodimerization domains, the near N-terminal domain (NND) and the twin-cysteine domain (TCD). The TCD crystal structure reveals a stable homodimer with a novel dimerization interface. While the NND homodimer is intrinsically less stable, using crosslinking mass spectrometry we show that within Mto1/2[bonsai] complexes, it can be reinforced by additional cooperative interactions involving both Mto2 and Mto1[bonsai]. We propose a model for Mto1/2[bonsai] complex architecture that is supported by functional analysis of mutants *in vivo*.

## INTRODUCTION

Microtubule (MT) nucleation by the *γ*-tubulin complex (*γ*-TuC) depends on a variety of accessory proteins that regulate *γ*-TuC localization, activity, and assembly state (Farache et al., 2018; Kollman et al., 2011; Lin et al., 2015; Paz and Luders, 2018; Petry and Vale, 2015; Roostalu and Surrey, 2017; Tovey and Conduit, 2018). The core element of the *γ*-TuC, the highly conserved heterotetrameric *γ*-tubulin small complex (*γ*-TuSC), contains two copies of *γ*-tubulin and one copy each of the related proteins GCP2 and GCP3 (in humans). In metazoan cells, multiple *γ*-TuSCs plus additional “non-core” *γ*-TuC proteins (e.g. GCP4, GCP5 and GCP6 in humans) assemble into a stable ∼2 MDa structure known as the *γ*-tubulin ring complex (*γ*-TuRC) (Oegema et al., 1999; Zheng et al., 1995). The *γ*-TuRC is thought to promote MT nucleation by binding multiple *αβ* tubulin dimers in an orientation that mimics a MT plus end, thereby overcoming kinetic barriers to nucleation at physiological tubulin concentrations. In addition to acting as a “template” for MT nucleation, the *γ*-TuRC may also stabilize spontaneous nucleation intermediates and MT minus ends (Anders and Sawin, 2011; Roostalu and Surrey, 2017; Wiese and Zheng, 2000). Interestingly, while non-core (i.e. *γ*-TuRC-specific) *γ*-TuC proteins appear to be critical for *γ*-TuRC assembly in metazoan cells (summarized in (Cota et al., 2017)), in some organisms (e.g. *Drosophila melanogaster, Aspergillus nidulans*, and fission yeast *Schizosaccharomyces pombe)*, these non-core proteins contribute only modestly to MT nucleation *in vivo* (Anders et al., 2006; Fujita et al., 2002; Venkatram et al., 2004; Verollet et al., 2006; Xiong and Oakley, 2009), and in other organisms (e.g. budding yeast *Saccharomyces cerevisiae*), such non-core proteins are completely absent (Lin et al., 2015; Tovey and Conduit, 2018). This diversity has suggested that in addition to the “classical” *γ*-TuRC of metazoan cells, there may be other mechanisms by which multiple *γ*-TuSCs can assemble to generate the functional equivalent of a *γ*-TuRC.

Proteins that contain a Centrosomin Motif 1 (CM1) domain (here referred to as “CM1 proteins”) regulate multiple aspects of *γ*-TuC function, including assembly into *γ*-TuRC-like structures (see below). The CM1 domain, a ∼60 amino-acid region that interacts with the *γ*-TuC, is conserved from yeasts to humans (Choi et al., 2010; Leong et al., 2019; Samejima et al., 2008; Sawin et al., 2004; Zhang and Megraw, 2007), and in most organisms CM1 proteins are either unique or present as a paralog pair. Characterized CM1 proteins include human CDK5RAP2/Cep215 and myomegalin/PDE4DIP, *Drosophila* centrosomin (Cnn), *Aspergillus* ApsB, budding yeast Spc110p, and fission yeast Mto1 and Pcp1 (summarized in (Lin et al., 2015; Tovey and Conduit, 2018)). Outside the CM1 domain, CM1 proteins are divergent in primary sequence; however, all CM1 proteins are ∼100 kDa or greater, and all contain the CM1 domain near their N-termini and multiple predicted coiled-coils in C-terminal regions (Suppl. Figure 1A). CM1 proteins were initially characterized for their role in recruiting the *γ*-TuC to MT organizing centers (MTOCs) such as centrosomes and yeast spindle pole bodies (SPBs) (Knop and Schiebel, 1997; Megraw et al., 2001; Sawin et al., 2004; Venkatram et al., 2004). More recently, *Drosophila* Cnn has been shown to dynamically assemble into large-scale scaffolds that, via their ability to recruit the *γ*-TuC, can modulate the MT-nucleation capacity of the centrosome during the cell cycle (Citron et al., 2018; Conduit et al., 2014; Feng et al., 2017). Importantly, independently of regulating localization of the *γ*-TuC, CM1 proteins from multiple organisms have also been shown to promote MT nucleation activity of the *γ*-TuC, in a range of *in vitro* and *in vivo* assays (Choi et al., 2010; Kollman et al., 2010; Leong et al., 2019; Lynch et al., 2014; Samejima et al., 2008). Thus, rather than serving as simple “adapters” that localize the *γ*-TuC to MTOCs, CM1 proteins may coordinate *γ*-TuC localization with *γ*-TuC activation, not only promoting *γ*-TuC activation at MTOC sites but also preventing undesired *γ*-TuC activation at non-MTOC sites.

In fission yeast, the two CM1 proteins Mto1 and Pcp1 have non-overlapping roles in regulating *γ*-TuC-dependent MT nucleation, providing an ideal system for analysis. Pcp1, which is essential for viability, is required for mitotic spindle formation within the nucleus (so-called “closed” mitosis) and is localized to the nucleoplasmic face of the SPB (Flory et al., 2002; Fong et al., 2010). By contrast, Mto1, which is nonessential, is required for all MT nucleation in the cytoplasm and is localized to multiple sites: the cytoplasmic face of the SPB, the cytoplasmic face of nuclear pore complexes, the lattice (sides) of pre-existing cytoplasmic MTs, and the cytokinetic actomyosin ring (Bao et al., 2018; Samejima et al., 2010; Sawin et al., 2004; Venkatram et al., 2004). (Although Mto1 is not essential for viability, it is required for normal polarized cell shape, nuclear positioning, and septum positioning.) Direct recruitment of Mto1 to the cytoplasmic face of the SPB appears to generate a sufficiently high local concentration of Mto1 to support its role in MT nucleation there (Samejima et al., 2010). However, to function at non-SPB sites, Mto1 requires a partner protein, Mto2 (Janson et al., 2005; Samejima et al., 2005; Venkatram et al., 2005). Mto2 is conserved in fungi in the Schizosaccharomyces and Pezizomycotina clades of Ascomycota and is predicted to contain several intrinsically disordered regions (Borek et al., 2015). *In vivo*, multiple Mto1 and Mto2 molecules associate to form a multimeric Mto1/2 complex that concentrates the proteins at MTOC sites (Lynch et al., 2014; Samejima et al., 2008; Samejima et al., 2010).

To date, little is known about how the Mto1/2 complex is assembled. However, a minimal active fragment of Mto1, Mto1[bonsai] (amino acid residues 131-549), has enabled initial investigations into Mto1/2 complex organization *in vivo* (Lynch et al., 2014). Mto1[bonsai] lacks N- and C-terminal residues (1-130 and 550-1115, respectively) that localize Mto1 to specific sites (Figure 1A, Suppl. Figure 1A), but it retains both the CM1 domain and the ability to bind Mto2. As a result, Mto1[bonsai] and Mto2 can form a multimeric Mto1/2[bonsai] complex that interacts with the *γ*-TuC. *In vivo*, Mto1/2[bonsai] complexes are present as freely-diffusing cytoplasmic puncta that also contain *γ*-TuSC proteins (Lynch et al., 2014). Puncta form equally well in the presence and absence of non-core *γ*-TuC proteins. Individual puncta nucleate single MTs in a spatially random manner, and fluorescence quantification of GFP-tagged Mto1[bonsai], Mto2, and *γ*-TuSC proteins within actively nucleating puncta has shown that they contain approximately 13 molecules each of Mto1[bonsai] and Mto2, and slightly more than half as many molecules each of *γ*-TuSC proteins Alp4 (GCP2 homolog) and Alp6 (GCP3 homolog) (Lynch et al., 2014). These features have suggested that each punctum may correspond to a *γ*-TuRC-like structure, generated without a requirement for non-core *γ*-TuC proteins.

**Figure 1.**
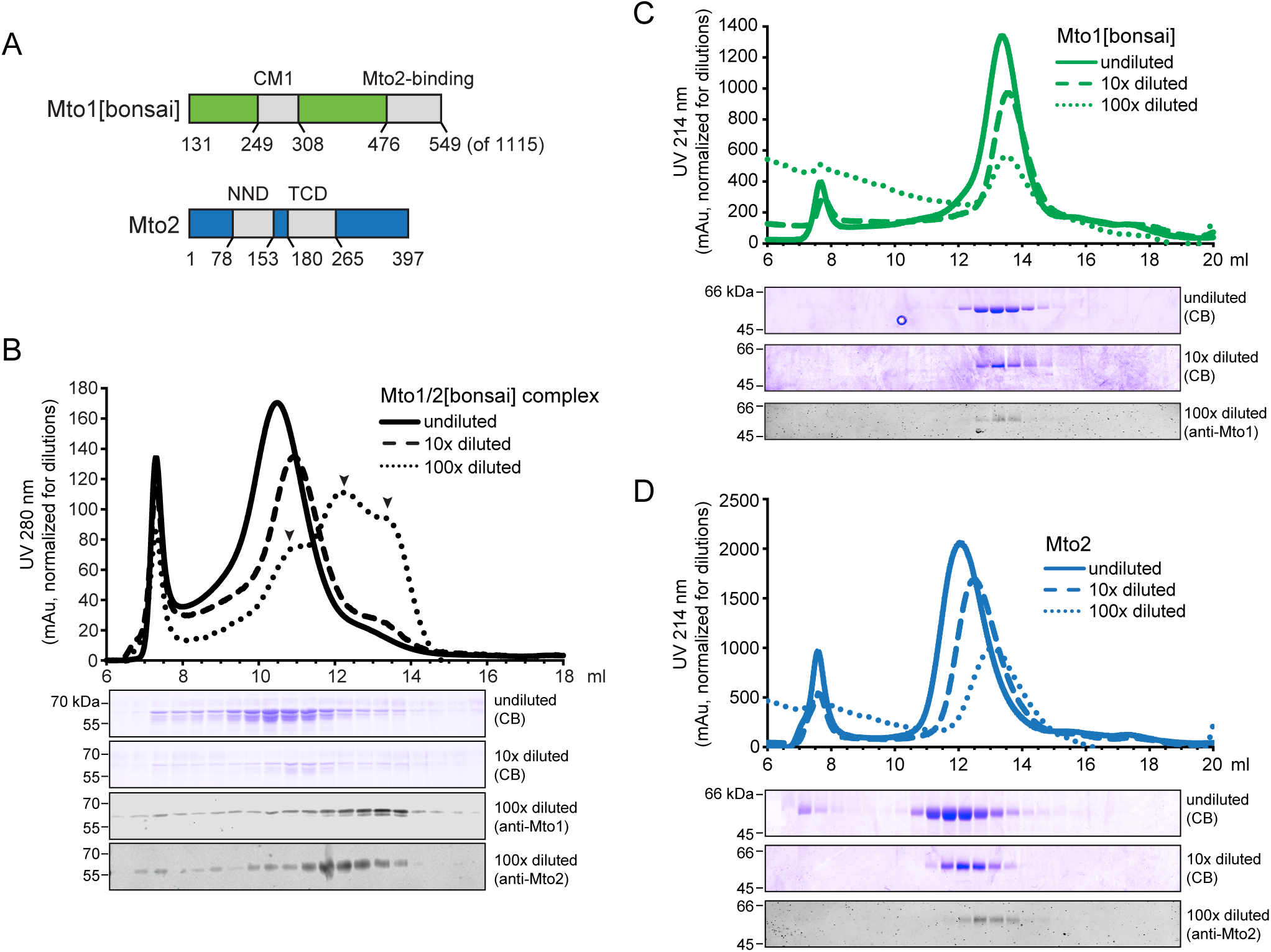
Mto1/2[bonsai] complex and Mto2 each form higher-older multimers *in vitro*. **(A)** Domain organization of Mto1[bonsai] (green) and Mto2 (blue). Residue numbering for Mto1[bonsai] corresponds to residues in full-length Mto1 (see Suppl. Figure 1A). CM1 indicates Centrosomin Motif 1. NND and TCD indicate Near-N-terminal Domain and Twin-Cysteine Domain. **(B)** Size-exclusion chromatography (SEC) of purified Mto1/2[bonsai] complex at different dilutions. Undiluted sample was injected at ∼9 mg/mL. Arrowheads indicate multiple peaks due to dissociation. Corresponding Coomassie Blue (CB)-stained SDS-PAGE and anti-Mto1 and anti-Mto2 western blots are shown underneath. In CB-stained gels, upper band is Mto1[bonsai], and lower band is Mto2. **(C, D)** SEC of purified Mto1[bonsai] (C) and purified Mto2 (D) at different dilutions. Undiluted samples were injected at ∼1.5 mg/mL (Mto1[bonsai]) and ∼2.5 mg/ml (Mto2). SDS-PAGE and western blots are shown underneath. See also Suppl. Figure 1.

Imaging of mutants *in vivo* has suggested that Mto1/2[bonsai] multimerization occurs independently of *γ*-TuSC and is critical for *γ*-TuSC multimerization into *γ*-TuRC-like structures. In mutants lacking Mto1/2[bonsai] puncta, *γ*-TuSC proteins lose punctate localization (Anders and Sawin, 2011; Lynch et al., 2014). By contrast, when the Mto1[bonsai] CM1 domain is mutated to prevent binding to *γ*-TuSC, some mutant Mto1/2[bonsai] complexes are still seen in puncta. Interestingly, Mto1[bonsai] loses punctate localization in *mto2Δ* deletion mutants, but some Mto2 is still seen in puncta in *mto1Δ* deletion mutants. Furthermore, overexpression of Mto2 increases puncta formation, while overexpression of Mto1[bonsai] does not. Collectively, these *in vivo* results have led to a model of hierarchical assembly in which Mto2 multimers drive the formation of Mto1/2[bonsai] multimers, which in turn help to assemble multiple *γ*-TuSCs into *γ*-TuRC-like structures (Lynch et al., 2014). At the same time, however, punctate localization of Mto2 is much less robust *in mto1Δ* cells compared to *mto1*[*bonsai*] cells, and punctate localization of Mto1/2[bonsai] is also less robust when Mto1/2[bonsai] cannot bind to *γ*-TuSCs (Lynch et al., 2014). Therefore, Mto1/2 multimerization, and the ability to generate *γ*-TuRC-like structures, may involve both hierarchical assembly and cooperative interactions involving all of the above proteins.

As the Mto1/2 complex simultaneously has key roles in *γ*-TuC localization, activation, and multimerization, a deeper understanding of how the multimeric Mto1/2 complex is assembled would provide critical insight into mechanisms of *γ*-TuC organization, particularly at non-centrosomal MTOCs, which are widespread in eukaryotic cells (Petry and Vale, 2015; Sanchez and Feldman, 2017; Wu and Akhmanova, 2017). In this work we investigate the architecture of the Mto1/2[bonsai] complex and the basis for its multimerization, through structural and biophysical analysis of purified proteins. We show that Mto1[bonsai] and Mto2 together are sufficient for formation of multimeric complexes *in vitro*, and that Mto2 alone can also multimerize. Analysis of Mto2 fragments further indicates that Mto2 multimerization depends on two independent homodimerization domains, and we solve the crystal structure of one of these domains. Using chemical crosslinking and mass spectrometry we provide evidence that additional interactions between Mto1[bonsai] and Mto2, and between Mto2 and itself, cooperate with Mto2 homodimerization domains to generate multimeric Mto1/2[bonsai] complexes. We validate these *in vitro* findings with experiments *in vivo*.

## RESULTS

### Both Mto1/2[bonsai] complex and Mto2 form higher-order multimers in vitro

We co-expressed Mto1[bonsai] and Mto2 in insect cells and co-purified them as an Mto1/2[bonsai] complex (Figure 1A, Suppl. Figure 1A,B). In size-exclusion chromatography (SEC), Mto1/2[bonsai] eluted as a single broad peak, with Mto1[bonsai] and Mto2 present in near-equal stoichiometry across the peak (Figure 1B, Suppl. Figure 1C). In a recent density-gradient centrifugation analysis, Mto1/2[bonsai] sedimented with a peak at 9-10S and a very long “tail”, extending to 20S and beyond, suggesting that the complex forms higher-order multimers (Leong et al., 2019). Consistent with this, using size-exclusion chromatography with multi-angle light scattering (SEC-MALS) we found that the Mto1/2[bonsai] SEC peak comprises a range of molecular masses (Suppl. Figure 1D, Table 1). While the average molecular mass within the peak was ∼250 kDa, a range of masses from ∼120 kDa to ∼1 MDa was seen at the shoulders of the peak. Dilution of Mto1/2[bonsai] shifted the SEC peak elution profile towards later-eluting species, indicating that multimerization is concentration-dependent (Figure 1B, “10X dilution”). At high dilution, the elution profile was shifted even further towards later-eluting species, but the Western-blot profiles of Mto1[bonsai] and Mto2 no longer matched, indicating that the two proteins had likely dissociated (Figure 1B, “100X dilution”). We could also reconstitute multimeric Mto1/2[bonsai] by mixing individually purified Mto1[bonsai] and Mto2; the complex generated by mixing had a similar elution profile to the complex generated from co-expression (Suppl. Figure 1E). Collectively, these results suggest that the recombinant Mto1/2[bonsai] complex exists as lower- and higher-order multimers in dynamic equilibrium *in vitro*.

**Table 1.** Native molecular mass of purified proteins, determined by SEC-MALS

| Protein/Complex/<br>Fragment | Input Protein<br>Concentration<br>(μM) | Theoretical<br>Mass (kDa) | Native Mass (kDa) | | Mw/<br>Theoretical<br>mass | Oligomer Status | Dispersity<br>(Mw/Mn) | Figure |
| --- | --- | --- | --- | --- | --- | --- | --- | --- |
|  |  |  | Mn | Mw |  |  |  |  |
| Mto1/2[bonsai] complex<br>(Mto1[bonsai] plus Mto2) | 45 | 94.4 | >1000-120 |  | N/A | Oligomer | N/A | Suppl. Fig. 1D |
| Mto1[bonsai] | 90 | 50.4 | 134.048 | 135.962 | 2.7 | Dimer | 1.014 | Suppl. Fig. 1F |
|  | 25 | 50.4 | 103.453 | 104.062 | 2.1 | Dimer | 1.006 | -- |
| Mto2 | 75 | 45.6 | >500-200 |  | N/A | Oligomer | N/A | -- |
|  | 30 | 45.6 | 500-80 |  | N/A | Oligomer | N/A | Suppl. Fig. 1G |
| Mto2-134A136A | 30 | 45.4 | 100.040 | 100.318 | 2.2 | Dimer | 1.003 | Suppl. Fig. 4D |
| Mto2-134AAA | 30 | 45.4 | 85.030 | 85.470 | 1.9 | Dimer | 1.005 | Suppl. Fig. 4E |
| Mto2 NND | 1800 | 8.6 | 20.677 | 21.087 | 2.5 | Dimer | 1.02 | Suppl. Fig. 2D<br>(and Fig. 2A)<br>(Mn and Mw<br>are for peak<br>region only) |
|  | 600 | 8.6 | 17.818 | 18.507 | 2.1 | Dimer | 1.039 |  |
|  | 180 | 8.6 | 12.915 | 13.273 | 1.5 | Dimer ~ Monomer | N/A |  |
|  | 90 | 8.6 | 11.856 | 12.787 | 1.5 | Dimer < Monomer | N/A |  |
| Mto2 NND-134A136A | 600 | 8.4 | 9.222 | 9.500 | 1.1 | Monomer | 1.030 | Suppl. Fig. 4B |
| Mto2 NND-134AAA | 150 | 8.4 | 8.226 | 8.565 | 1.0 | Monomer | 1.041 | Suppl. Fig. 4C |
| Mto2 TCD | 1000 | 9.3 | 17.469 | 17.549 | 1.9 | Dimer | 1.005 | Suppl. Fig. 2E<br>(and Fig. 2D) |
|  | 500 | 9.3 | 16.524 | 16.571 | 1.8 | Dimer | 1.003 |  |
|  | 100 | 9.3 | 17.049 | 17.093 | 1.8 | Dimer | 1.003 |  |

Mn, number-averaged molecular mass; Mw, weight-averaged molecular mass.
For simplicity, names of proteins do not include 6x-His tag or linker/cleavage sequences, although these are included in mass calculations.

To determine what drives Mto1/2[bonsai] multimerization, we characterized Mto1[bonsai] and Mto2 individually by SEC-MALS and SEC-dilution analysis. In SEC-MALS, Mto1[bonsai] alone was dimeric, while Mto2 alone showed a range of molecular masses (estimated ∼80-500 kDa), suggesting multimerization similar to that observed for Mto1/2[bonsai] complex (Suppl. Figure 1 F,G; Table 1). In SEC-dilution analysis, dilution did not significantly change the Mto1[bonsai] elution profile, indicating that the Mto1[bonsai] dimer is stable (Figure 1C). By contrast, dilution shifted the Mto2 elution profile towards later-eluting species (Figure 1D). These *in vitro* results suggest that concentration-dependent multimerization of the Mto1/2[bonsai] complex is driven largely by the Mto2 component. This is consistent with *in vivo* studies showing that Mto2 is present in puncta independently of Mto1 (albeit more weakly than when Mto1[bonsai] is present), and that Mto2 from yeast cell lysates has a higher sedimentation coefficient when concentrated (Lynch et al., 2014).

### Mto2 contains two independent homodimerization domains

We next investigated the basis for Mto2 multimerization. Although a large proportion of Mto2 is predicted to be intrinsically disordered (Borek et al., 2015), the two most conserved regions of Mto2 have significant predicted alpha-helical content (Figure 1A, Figure 2). We therefore purified and characterized these two regions individually (Suppl. Figure 2A).

**Figure 2.**
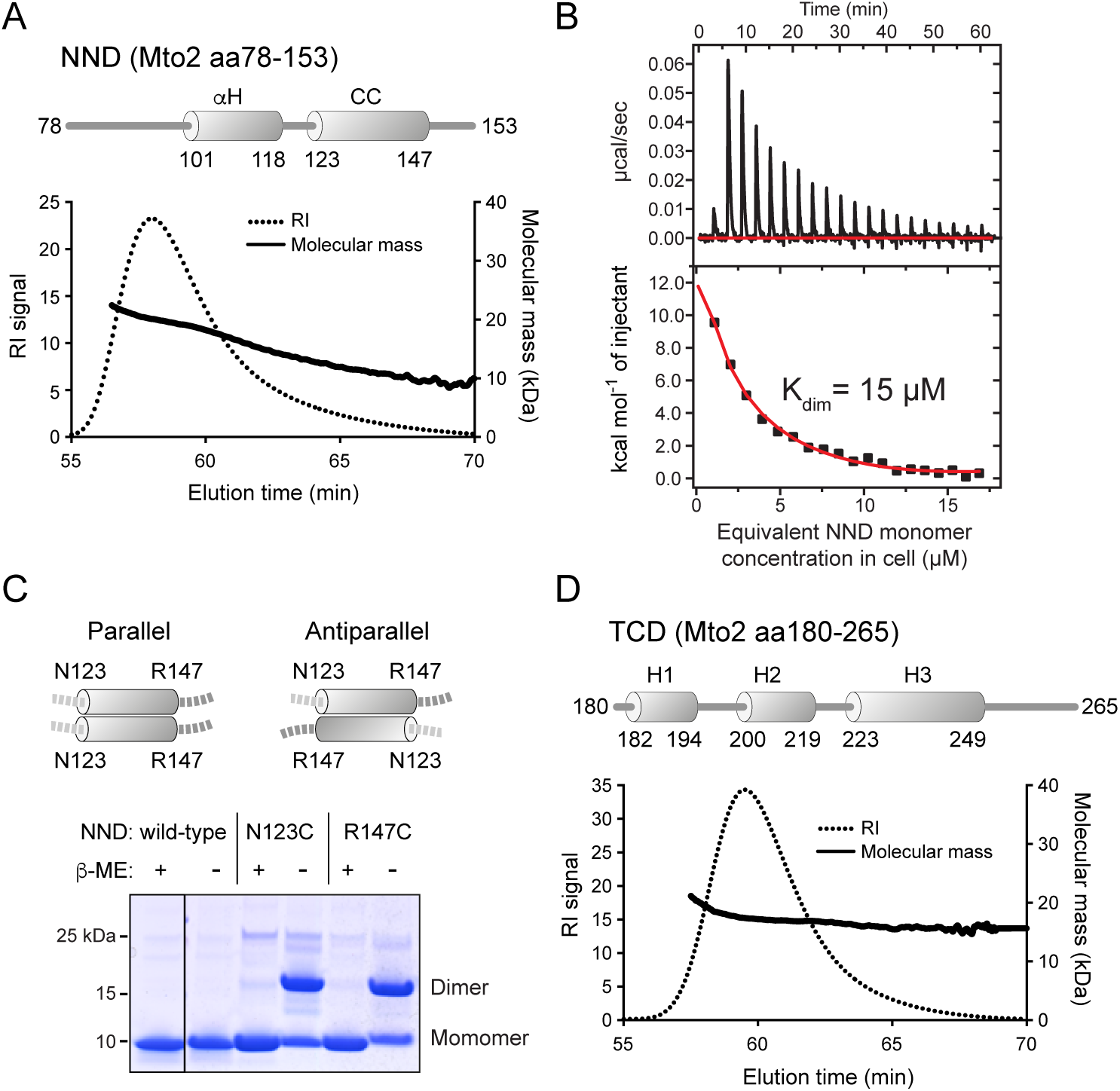
The Mto2 Near-N-terminal Domain (NND) and Twin-Cysteine Domain (TCD) each form homodimers. **(A)** Size-exclusion chromatography with multi-angle laser-light scattering (SEC-MALS) analysis of the NND. Sample was injected at ∼5 mg/mL. Schematic of predicted alpha-helices indicates non-coiled-coil (*α*H) and coiled-coil (CC) regions. In the tail following the protein peak, gradual decrease in molecular mass (to 50% of peak value) indicates dissociation of dimers into monomers. See also Suppl. Figure 2B,D. **(B)** Isothermal titration calorimetry analysis of NND homodimer dissociation. **(C)** SDS-PAGE of wild-type NND and mutants NND-N123C and NND-R147C, under reducing and non-reducing conditions (with and without *β*-mercaptoethanol; *β*-ME). Dimer bands indicate intermolecular disulfide bond formation under non-reducing conditions, suggesting that CC region forms a parallel homodimer. **(D)** SEC-MALS analysis of the TCD. Sample was injected at ∼10 mg/mL. Schematic indicates predicted alpha helices (H1, H2, H3). In the tail following the protein peak, molecular mass remains constant. See also Suppl. Figure 2C,E.

The first region (residues 78-153) contains a predicted alpha helix (residues 101-118) followed by a short, predicted alpha-helical coiled-coil (residues 123-147); we refer to this region as the NND (“Near-N-terminal domain”; Figure 1A, Figure 2A). SEC-MALS and SEC-dilution analysis showed that the NND alone forms homodimers that dissociate upon dilution (Figure 2 A, Suppl. Figure 2B,D; Table 1). Consistent with this, isothermal titration calorimetry (ITC) revealed an NND homodimer dissociation constant (K_d_) of 15 µM (Figure 2B). We conclude that the NND alone forms a weak homodimer.

The predicted coiled-coil within the NND led us to test whether it forms a parallel or antiparallel homodimer. We generated mutant NND proteins containing a cysteine residue either at the beginning or the end of the coiled-coil (mutants NND-N123C and NND-R147C, respectively; Figure 2C). We reasoned that if NND-N123C forms a parallel homodimer, then cysteine 123 from one monomer within the dimer should be able to form a disulfide bond with cysteine 123 from the other monomer. By contrast, if NND-N123C forms a antiparallel homodimer, such a disulfide bond would not form. Analogous arguments apply to cysteine 147 in an NND-R147C homodimer. Consistent with a parallel orientation, both NND-N123C and NND-R147C migrated as dimers on non-reducing SDS-PAGE but as monomers on reducing SDS-PAGE (Figure 2C). This suggests that the wild-type homodimer is parallel.

The second predicted alpha-helical region of Mto2 (residues 180-265) contains three alpha helices; because the third helix contains two highly-conserved cysteine residues (Borek et al., 2015), we refer to this region as the TCD (“Twin-Cysteine Domain”; Figure 1A, Figure 2D). SEC-MALS indicated that, like the NND, the TCD forms homodimers (Figure 2D, Suppl. Figure 2E, Table 1). However, unlike the NND, the TCD did not dissociate upon dilution (Suppl. Fig 2C,E). Thermal denaturation assays further showed that the TCD has a high melting temperature (∼78°C; Suppl. Figure 2F). We conclude that the TCD forms a particularly stable homodimer.

### The Mto2 Twin-Cysteine Domain (TCD) homodimerizes via a novel hydrophobic interface

To investigate the basis for homodimerization of the TCD, we determined its structure by X-ray crystallography (Figure 3, Table 2). Phases were obtained using single-wavelength anomalous dispersion from selenomethionine-labelled protein crystals that diffracted to 1.9 Å in the space group *P*2_1_2_1_2_1_. After model building and refinement in this space group, the model was used to determine phases by molecular replacement in a native dataset that diffracted to 1.5 Å in space group *P*6_1_. Residues 180-250 (chain A) and 180-248 (chain B) could be built into the electron density map. Each TCD monomer forms a three-helix bundle, in which the third helix is the longest and extends beyond the core (Figure 3A). Within each monomer, the three helices are held together by extensive hydrophobic interactions (Suppl. Figure 3); these involve several leucine and valine side chains, as well as the two conserved cysteine residues, which, although polar, have a strong hydrophobic character (Nagano et al., 1999). The two monomers dimerize via a similarly hydrophobic interface (Suppl. Figure 3), burying a surface area (per monomer) of 974.6 Å^2^, and a comparison of the buried dimer interface with the solvent-exposed surface revealed a distinct segregation of non-polar and polar surfaces (Figure 3B). The large proportion of buried surface area of the TCD dimer and its hydrophobic nature explain why it is not easily disrupted by dilution (Lo Conte et al., 1999) and is essentially an obligate, non-dissociable homodimer.

**Figure 3.**
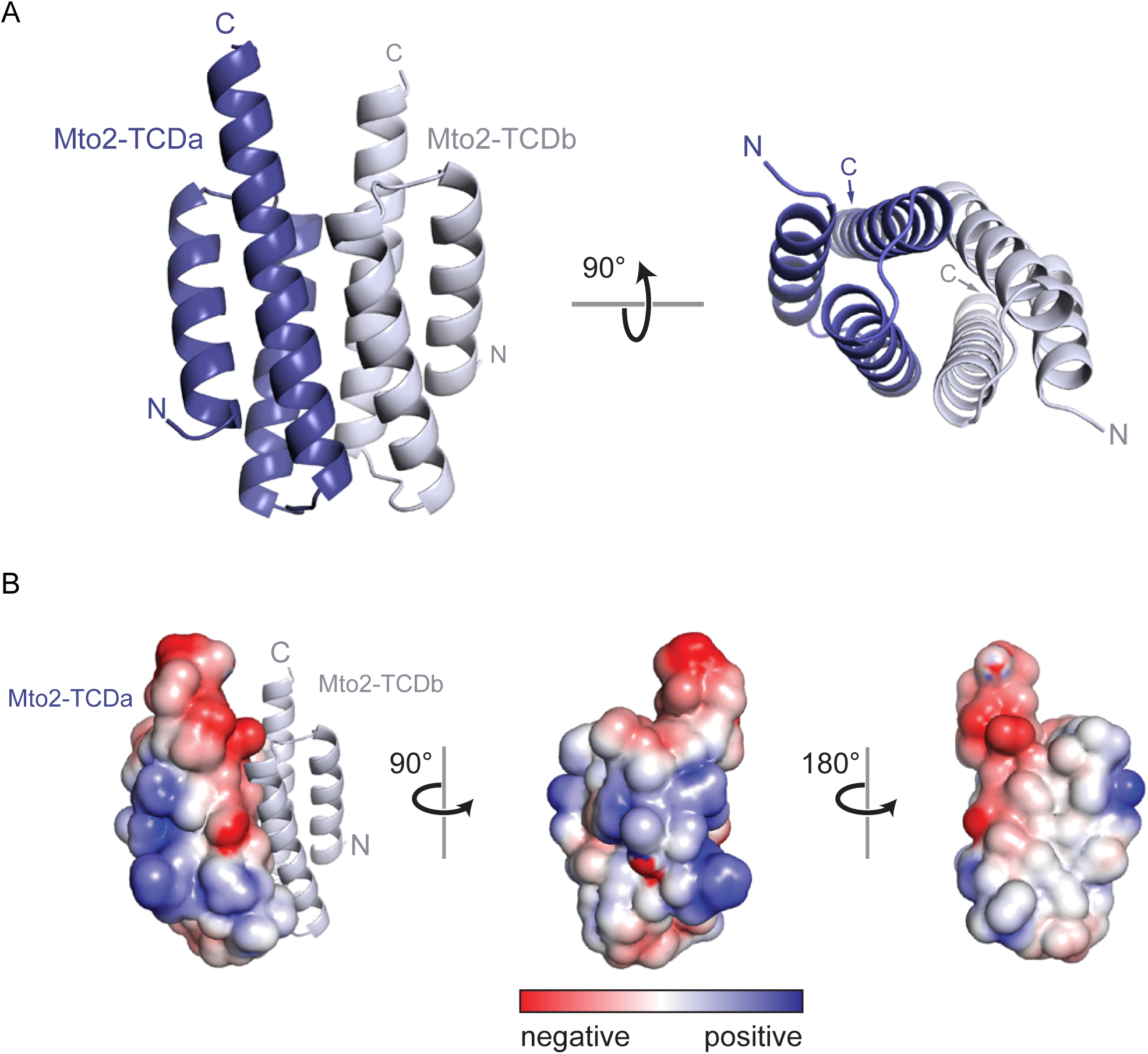
X-ray crystal structure of the Mto2 Twin-Cysteine Domain (TCD). **(A)** Ribbon diagram of the TCD dimer. The two monomer chains (TCDa and TCDb) are in different shades of blue. Rotated view shows proximity of the two three-helix bundles. See also Suppl. Figure 3. **(B)** Surface-charge representation of the TCD monomer. Left panel includes ribbon diagram of second monomer, to show orientation of the dimer relative to A. Note absence of charged residues at the hydrophobic dimer interface (right panel, white surface).

**Table 2.** Data collection and refinement statistics

|  | Se-Methionine SAD | Native (6GDJ) |
| --- | --- | --- |
| Beamline | DLS-I04 | DLS-I02 |
| Wavelength (Å) | 0.979 | 1.0 |
| Space group | P2<sub>1</sub>2<sub>1</sub>2<sub>1</sub> | P6<sub>1</sub> |
| Unit cell | a=54.62Å b=73.86Å, c=98.43Å; $\alpha=\beta=\gamma=90^\circ$ | a=b=73.7, c=54.4Å $\alpha=\beta=90^\circ$ $\gamma=120^\circ$ |
| Resolution (High resolution shell) | 24.8-1.9 (2.0-1.9) Å | 41.4-1.5 (1.58-1.50) Å |
| Unique reflections | 57043 | 26976 |
| Completeness (%) |  | 99.9 (99.6) |
| Anomalous Completeness (%) | 94.2 (71.2) |  |
| Multiplicity | 3.2 | 5.1(4.9) |
| I/s(I) | 12.6 (3.5) | 6.6 (2.6) |
| Rmerge | 0.081 (0.268) | 0.155(0.455) |
| CC(1/2) | 99.6 (93.1) | 97.4 (92.4) |
| Refinement: |  | CCP4 Refmac5 |
| R<sub>free</sub> |  | 0.183 |
| R<sub>work</sub> |  | 0.152 |
| Total number of protein atoms |  | 1033 |
| Total number of non-protein atoms |  | 68 |
| Average B-factor (protein atoms) |  | 23.51 |
| RMSD Bond Length (Å) |  | 0.020 |
| RMSD Bond Angles (Degree) |  | 1.804 |
| RMSD Chiral Centre |  | 0.128 |
| Ramachandran plot (COOT validation) |  |  |
| In preferred Regions |  | 100% (137) |
| In allowed Regions |  | 0.00% (0) |
| Outliers |  | 0.00% (0) |

To determine the relatedness of this structure to other proteins, we queried the TCD against known structures using PDBeFOLD (Krissinel and Henrick, 2004). The closest structural relatives are the Staphylococcal complement inhibitor D (root mean square deviation, rmsd=1.7 Å) and acetyl co-A binding proteins from human, rice and cattle (rmsd=2.0-2.1 Å) (Garcia et al., 2012; Guo et al., 2017; Taskinen et al., 2007; van Aalten et al., 2001). However, these proteins are monomeric, and the acetyl co-A binding proteins have an additional helix that binds against the equivalent of the surface that is buried in the TCD homodimer. The Mto2 TCD thus appears to use a novel interface for stable homodimerization.

### Both Mto2 homodimerization domains contribute to Mto2 multimerization

Our finding that Mto2 contains two independent homodimerization domains suggested a general model for Mto2 multimerization in which one Mto2 molecule can interact with another Mto2 molecule via NND dimerization and with a different Mto2 molecule via TCD dimerization. Designating NND dimerization as an “A-A” interaction and TCD dimerization as a “B-B” interaction, we refer to this model for Mto2 multimerization as the “ABBA” model (Figure 4A).

**Figure 4.**
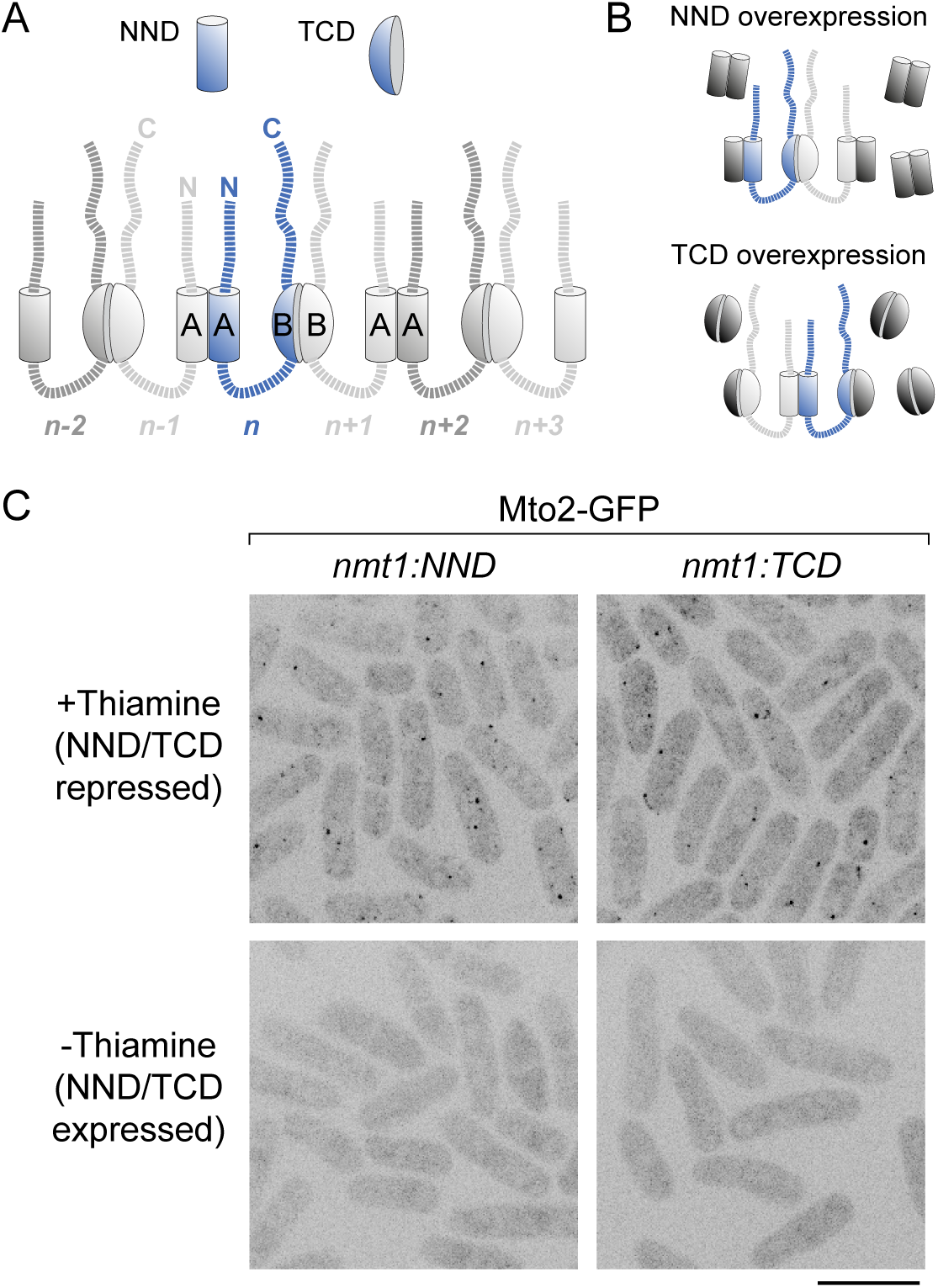
Both the Mto2 NND and TCD contribute to Mto2 puncta formation *in vivo*. **(A)** “ABBA” model for Mto2 multimerization. Mto2 polypeptides are linked by alternating NND-NND (“A-A”) and TCD-TCD (“B-B”) interfaces. The number of polypeptides shown is arbitrary and could be extended in either direction (see Discussion). **(B)** Schematic of how NND or TCD overexpression could disrupt Mto2 multimerization, according to the ABBA model. **(C)** Mto2-GFP puncta in *mto1Δ* cells when NND or TCD transgenes are repressed (top panels), and absence of Mto2-GFP puncta when NND or TCD transgenes are overexpressed (bottom panels). Scale bar, 10 µm. See also Suppl. Figure 4.

The ABBA model predicts that both homodimerization domains should be important for Mto2 multimerization. To test the role of the NND in multimerization, we mutated amino-acid residues predicted to stabilize the coiled-coil (Suppl. Figure 4A), to make mutant NND proteins NND-134A136A and NND-134AAA. SEC-MALS showed that in the context of the NND alone, the mutant proteins formed monomers rather than dimers (Suppl. Figure 4B,C, Table 1). Similarly, when the same mutations were introduced into full-length Mto2, the mutant proteins failed to form multimers and instead formed dimers, presumably via the TCD-TCD dimer interface (Suppl. Figure 4D,E, Table 1). Overall, these results suggest that NND dimerization is important for Mto2 multimerization (because we subsequently found that the NND also interacts with Mto1, *in vivo* phenotypes are described later below)

In parallel with these experiments, we attempted to construct analogous mutants to “monomerize” the TCD. However, all mutations tested led to unfolding of the TCD. It is likely that disrupting the dimer interface exposes a large, hydrophobic surface that destabilizes the fold in an aqueous environment (see Figure 3; Suppl. Figure 3).

According to the ABBA model, overexpression of either the NND or the TCD might have a dominant-negative effect on formation of Mto2 puncta *in vivo*, because either domain could potentially dimerize with endogenous full-length Mto2 to terminate ABBA-based multimerization (Figure 4B). To test this, we overexpressed the NND and TCD (separately) in cells expressing endogenous levels of Mto2-GFP (Lynch et al., 2014). To avoid any confounding effects due to cooperative interactions with Mto1 or *γ*-TuSC, we used *mto1Δ* cells. Overexpression of either the NND or TCD led to disappearance of Mto2-GFP puncta, leaving only cytosolic fluorescence (Figure 4C, Suppl. Figure 4F). These results suggest that Mto2 multimerization requires both NND-NND and TCD-TCD interactions, consistent with the ABBA model.

### Additional Mto2-Mto2 interactions stabilize Mto2 multimerization

While the ABBA model provides an attractive explanation for Mto2 multimerization, the model—at least in its simplest form—presented two potential issues. First, while the K_d_ for NND homodimerization *in vitro* is 15 µM, we determined that the concentration of endogenous Mto2 in the fission yeast cytosol is ∼160 nM, approximately 100 times lower than the K_d_ (Suppl. Figure 5A). As a result, the NND-NND dimer interface would not be expected to be stable *in vivo* unless supported by additional interactions. Second, if Mto2 were simply to consist of two dimerization domains connected by flexible disordered linkers, then two individual Mto2 molecules might be expected to form a “closed bivalent dimer” rather than participate in higher-order multimerization (Suppl. Figure 5B). As this is not observed, it suggested that Mto2 multimerization may involve additional structural features that prevent closed bivalent dimer formation.

To investigate these questions, we analyzed Mto2 by crosslinking mass spectrometry (CLMS) (Leitner et al., 2016; O’Reilly and Rappsilber, 2018; Sinz, 2018), using the zero-length crosslinker 1-ethyl-3-(3-dimethylaminopropyl)carbodiimide (EDC). We observed several intermolecular Mto2-Mto2 crosslinks that support the parallel nature of the NND dimer and dimerization of the TCD (Figure 5A,B). More importantly, multiple crosslinks indicated Mto2-Mto2 interactions extending beyond the homodimerization domains (Figure 5A). In particular, many residues in the Mto2 predicted disordered N- and C-terminal regions could be crosslinked to a central portion of the NND (residues 103-131).

**Figure 5.**
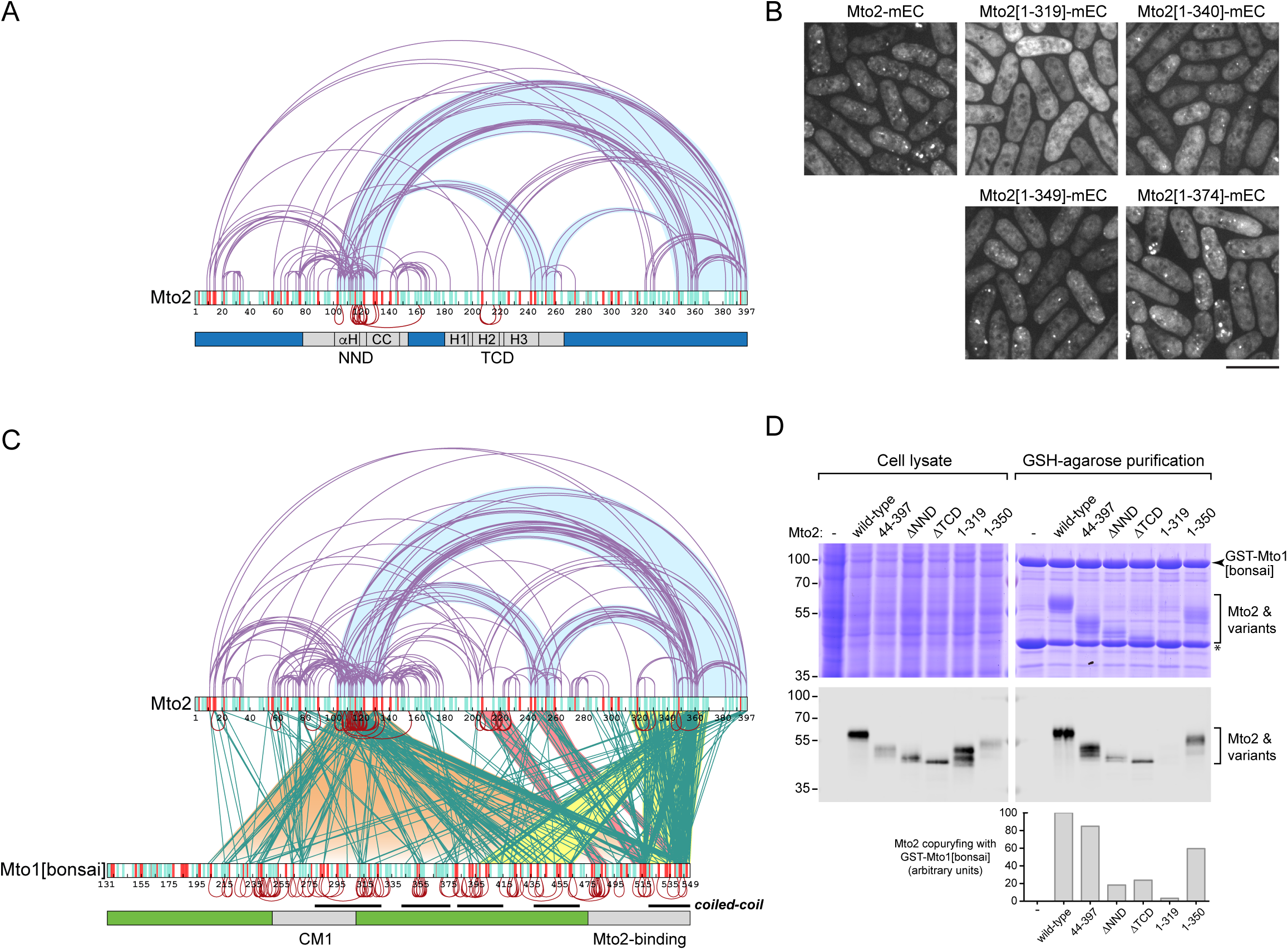
Mto2-Mto2 and Mto2-Mto1 interactions identified by crosslinking mass spectrometry. **(A)** Mto2-Mto2 interactions identified by EDC crosslinking of Mto2 alone. Light-blue shading indicates links between C terminus and NND, and links from both of these regions to TCD. **(B)** *In vivo* puncta formation for wild-type Mto2-mECitrine (Mto2-mEC) and Mto2 C-terminal truncations, in *mto1Δ* strain background. No puncta are seen with truncation Mto2[1-319]-mEC. **(C)** Mto1-Mto2, Mto2-Mto2, and Mto1-Mto1 interactions identified by EDC crosslinking of Mto1/2[bonsai] complex. For clarity, for Mto1-Mto1 crosslinks, only unambiguously intermolecular crosslinks are shown. Additional Mto1-Mto1 crosslinks from the same experiment are shown in Suppl. Figure 5C. **(D)** Copurification of wild-type Mto2 and Mto2 mutants with GST-Mto1[bonsai]. Upper panels show SDS-PAGE Coomassie Blue stain. Lower panels show anti-Mto2 western blot, with quantification. In A and C, red and green links are intermolecular; purple links could in principle be either intra- or intermolecular (see Discussion). Within each protein, potentially crosslinkable residues are indicated by green and red vertical lines, representing nucleophilic and carboxylic acid groups, respectively.

To test if these regions contribute to Mto2 multimer stabilization *in vivo*, we replaced endogenous Mto2 (in *mto1Δ* cells) with mildly overexpressed Mto2 N- and C-terminal truncations tagged with mECitrine. All N-terminal truncations were observed in puncta similar to those seen with full-length Mto2 (Suppl. Figure 5C), indicating that the Mto2 N-terminus (i.e. residues 1-89) is not required for punctate localization. Among C-terminal truncations (Figure 5B), Mto2[1-349]-mEcitrine and more distal C-terminal truncations were also observed in puncta. Strikingly, however, Mto2[1-319]-mEcitrine showed no punctate localization, while Mto2[1-340]) showed some punctate localization. These results suggest that a region of Mto2 near the C-terminus (approximately, residues 320-350) helps to stabilize Mto2 multimers *in vivo*, most likely by supporting the NND-NND interaction (see Discussion).

We also identified crosslinks from the NND to a region near the end of Helix 3 in the TCD, and from the TCD to the C-terminal region (Figure 5A). Together with the results described above, this suggests that the NND, the C-terminal region and a portion of the TCD may all be in close proximity; if so, this could potentially constrain the orientation of the TCD relative to the NND in a manner that prevents formation of a closed bivalent dimer (see Discussion).

### Multiple regions of Mto2 contribute to interaction with Mto1

Because previous work suggested that Mto1 binding to Mto2 may cooperatively stabilize Mto1/2[bonsai] puncta *in vivo* (Lynch et al., 2014), we used EDC and CLMS to analyze Mto1-Mto2 interactions within the Mto1/2[bonsai] complex. Mto1 amino-acid residues 476-549 have been shown to be required for binding to Mto2 (Samejima et al., 2005), but the regions of Mto2 involved in binding to Mto1 have not yet been investigated.

Most crosslinks from Mto2 to Mto1[bonsai] could be grouped into two major classes (Figure 5C). The first class (Figure 5C, yellow shading) includes crosslinks from the Mto2 C-terminal region (in particular, residues 317-369) to the Mto1 region previously implicated in Mto2 binding (in particular, residues 516-549), as well as to Mto1 residues 398-464. The second class (Figure 5C, orange shading) includes crosslinks from a portion of the Mto2 NND (residues 96-131) to most of the length of Mto1[bonsai] (residues 203-549), including the region previously implicated in Mto2 binding. In addition to these two classes, a smaller number of crosslinks were observed from Helix 2 of the Mto2 TCD (Mto2 residues 204-223) to Mto1 residues 479-489, and from a region near the end of Helix 3 of the TCD (Mto2 residues 242-257) to Mto1 residues 509-517 and 540-547 (Figure 5C, red shading).

Based on these results, we next tested which regions of Mto2 are important for interaction with Mto1[bonsai]. We expressed Mto2 truncation and internal-deletion mutants together with GST-Mto1[bonsai] in insect cells and assayed Mto2 binding to GST-Mto1[bonsai] through copurification on glutathione-agarose (Figure 5D). N-terminal truncation Mto2[44-397] bound to GST-Mto1[bonsai] similarly to wild-type Mto2, while internal deletions Mto2-ΔNND and Mto2-ΔTCD showed strongly decreased binding. Interestingly, C-terminal truncation Mto2[1-319], which contains both the NND and the TCD, showed almost no binding to GST-Mto1[bonsai]. Binding of a more distal C-terminal truncation, Mto2[1-350], was much better than Mto2-ΔNND, Mto2-ΔTCD, and Mto2[1-319], although not as good as wild-type Mto2. Collectively, these results indicate that multiple regions of Mto2—the NND, the TCD, and a C-terminal region (residues 320-350)— contribute to interaction with Mto1. As our CLMS data suggest that these different regions may all be in close proximity, this supports the idea that the Mto1/2[bonsai] complex may be stabilized by cooperative interactions involving some or all of these regions. This view is further supported by the fact that the C-terminal region of Mto2 required for Mto1 binding (i.e. residues 320-350) is the same region that is required for Mto2 punctate localization (Figure 5B).

In Mto1/2[bonsai] CLMS experiments, Mto2-Mto2 crosslinks were similar to those seen in experiments involving Mto2 alone, suggesting that Mto2 is unlikely to undergo major structural changes upon binding to Mto1 (Figure 5A, Figure 5C). The CLMS experiments also revealed multiple short-range intermolecular Mto1-Mto1 crosslinks in predicted coiled-coil regions (Figure 5C). Together with Mto1[bonsai] SEC-MALS and SEC-dilution data (Figure 1C, Suppl. Figure 1F, Table 1), this strongly suggests that Mto1[bonsai] is a parallel coiled-coil dimer. Mto1[bonsai] also showed longer-range crosslinks with itself, suggesting that the parallel coiled-coil dimer may fold back on itself (Suppl. Figure 5D); however, this was not investigated further.

### Mutations in the Mto2 NND and TCD regions phenocopy Mto2 deletion

Our results thus far suggest that the Mto2 NND plays a critical role in Mto2 function, as it not only homodimerizes but also interacts with other regions of Mto2 and with Mto1[bonsai]; all of these interactions may contribute to stabilization of Mto1/2[bonsai] multimers. To test the role of the NND *in vivo*, we replaced wild-type *mto2+* with *mto2* mutants containing charged-to-alanine mutations within the NND region (Suppl. Figure 6A; this includes the *mto2-134AAA* mutant characterized *in vitro*). We used strains expressing Mto1[bonsai]-GFP and mCherry-tubulin and assayed mutants for punctate Mto1[bonsai]-GFP localization, microtubule organization, and coimmunoprecipitation of Mto2 and *γ*-tubulin with Mto1[bonsai]-GFP. All mutant Mto2 proteins were stably expressed at near wild-type levels, although some mutants showed slightly altered migration patterns on SDS-PAGE (Figure 6B), probably because of small changes in the spectrum of phosphorylated isoforms (Borek et al., 2015). In *mto2-134AAA* and *mto2-129AAA* mutants, which contain mutations in the NND coiled-coil (Supp. Figure 4A; Suppl. Figure 6A), very few Mto1[bonsai]-GFP puncta were observed, and MT morphology and bundle number were highly aberrant compared to wild-type (*mto2+)* cells. These phenotypes were similar to those of *mto2Δ* mutants (Figure 6A; Suppl. Figure 6B,C), although Mto1[bonsai]-GFP puncta were slightly more numerous in *mto2-134AAA* and *mto2-129AAA* compared to *mto2Δ*. Neither Mto2-134AAA nor Mto2-129AAA was efficiently coimmunoprecipitated with Mto1[bonsai]-GFP, and in these mutants *γ*-tubulin was also poorly coimmunoprecipitated with Mto1[bonsai]-GFP (Figure 6B-D). The double mutant *mto2-129AAA134AAA* had a similar (i.e. non-additive) phenotype. These results suggest that *mto2-134AAA* and *mto2-129AAA* are nearly complete loss-of-function mutants *in vivo*. By contrast, mutants *mto2-103AAA* and *mto2-112AA116AA*, which contain mutations in the non-coiled-coil portion of the NND, showed essentially wild-type phenotypes, indicating that not all mutations in the NND are deleterious.

**Figure 6.**
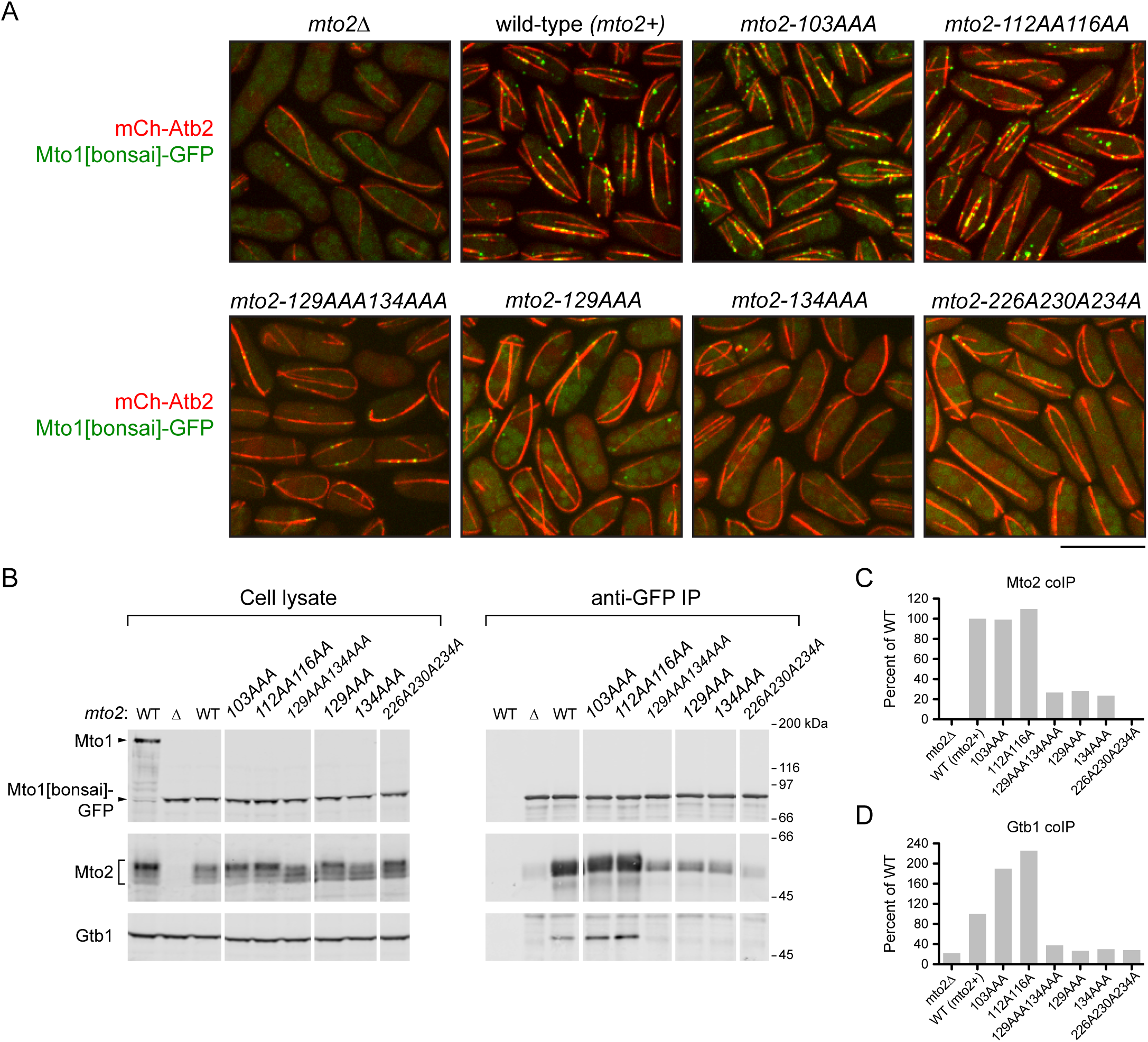
Mutations in the Mto2 NND and TCD impair Mto1/2 function *in vivo*. **(A)** Microtubule organization and Mto1[bonsai]-GFP puncta formation in wild-type cells (*mto2+)* and in the *mto2* mutants indicated. Quantification of microtubule organization is shown in Suppl. Figure 6. Scale bar, 10 µm. **(B)** Western blots for Mto1, Mto2, and *γ*-tubulin (Gtb1), showing coimmunoprecipitation with Mto1[bonsai]-GFP in wild-type cells and in the *mto2* mutants indicated. Left-most lane is from cells with untagged full-length Mto1. **(C, D)** Quantification of coimmunoprecipitation of Mto2 (C) and Gtb1 (D) with Mto1[bonsai]-GFP, from blots in B.

In parallel with these experiments, we generated a mutant, *mto2-226A230A234A*, containing charged-to-alanine mutations within Helix 3 of the TCD, close to the Mto2 residues that could be crosslinked to Mto1(Suppl. Figure 6A; Figure 5C). Like *mto2-134AAA* and *mto2-129AAA* cells, mutant *mto2-226A230A234A* cells showed strongly aberrant MT organization and impaired Mto1[bonsai]-GFP puncta formation (Figure 6A; Suppl. Figure 6B,C). Consistent with this, neither Mto2-226A230A234A nor *γ*-tubulin was coimmunoprecipitated with Mto1[bonsai]-GFP from these cells (Figure 6B-D). These results indicate that, like the NND, the TCD is critical for Mto2 function *in vivo*.

## DISCUSSION

In this work we provide insights into assembly and architecture of the Mto1/2 microtubule nucleation complex, through biophysical and structural analysis of the Mto1/2[bonsai] complex, a freely-diffusible, minimal form of the Mto1/2 complex that retains Mto1/2’s ability to promote *γ*-tubulin complex-dependent microtubule nucleation (Leong et al., 2019; Lynch et al., 2014). Previous imaging of Mto1/2[bonsai] puncta *in vivo* suggested that higher-order structure and stability of Mto1/2[bonsai] complex may depend on underlying multimerization of Mto2, as well as additional cooperative interactions (Lynch et al., 2014). Here we reproduce this multimerization *in vitro* and demonstrate how it can be achieved at a structural level by the Mto2 NND and TCD homodimerization domains (i.e. the ABBA model). In addition, we show that additional Mto2-Mto2 and Mto1-Mto2 interactions can cooperatively reinforce multimer stability by binding to the NND, which appears to serve as a hub for Mto1/2 function. Collectively, these results, together with analysis of mutants *in vivo*, suggest a model in which multiple inter- and intramolecular interactions stabilize Mto1/2[bonsai] architecture (Figure 7, Suppl. Figure 7).

**Figure 7.**
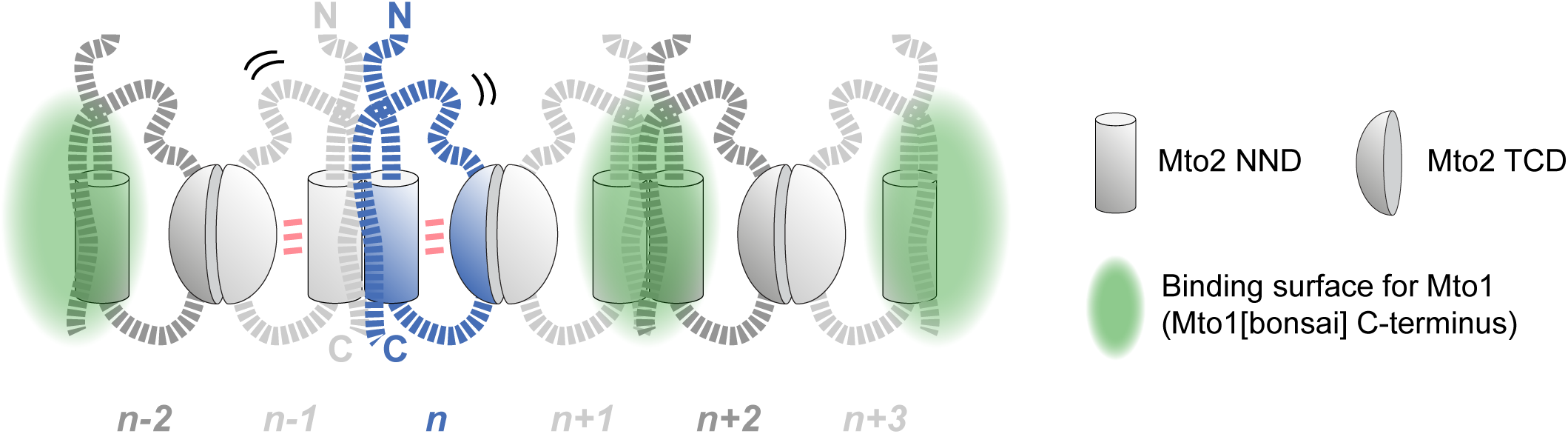
Model for architecture of Mto1/2 complex. The model builds on the ABBA model of Figure 4, incorporating additional Mto2-Mto2 and Mto2-Mto1 interactions identified by CLMS and truncation/deletion analysis. To illustrate Mto2-Mto2 interactions more clearly, the binding site for Mto1 is not shown at the interface between Mto2*_n_* and Mto2*_n-1_*. Pink triple bars indicate that the NND and TCD are close enough in space to allow EDC-dependent crosslinks. Squiggles represent mobility of predicted disordered regions, which may confer flexibility on the larger complex. Model is schematic and does not try to include more detailed elements. See also Suppl. Figure 7.

We used CLMS to identify Mto2-Mto2 and Mto1-Mto2 interactions beyond the Mto2 NND and TCD homodimerization domains. A potential issue with CLMS is that the positions of individual crosslinks may not correlate precisely with functionally important interactions. One reason for this is that only a subset of amino-acid side chains are compatible with crosslinking chemistry; another is that the side chains most intimately involved in interaction may be buried and poorly accessible to crosslinkers. In addition, if Mto1/2[bonsai] has a flexible structure (which seems likely, given Mto2’s predicted disordered regions), then CLMS results may represent the superposition of several different conformations rather than a single conformation. However, in spite of these caveats, groups of neighboring crosslinks should indicate general regions of interaction that can be further tested by mutational analysis. This approach allowed us to determine, for example, that Mto2 residues 320-350 are particularly important both for Mto1[bonsai]-Mto2 interaction *in vitro* and also for Mto2 puncta *in vivo*.

Another potential issue associated with CLMS concerns whether Mto2-Mto2 crosslinks are inter- or intramolecular. When crosslinks are between two identical or overlapping peptides from the same protein species, they are unambiguously intermolecular (e.g. the crosslinks along the length of the Mto1[bonsai] coiled-coil dimer in Figure 5C). However, when crosslinks are between non-overlapping peptides from the same protein and there are two or more copies of that protein within the complex being studied, it is generally not possible to distinguish between inter- and intramolecular interactions. This might seem to complicate the interpretation of Mto2 CLMS data, because the longer-range Mto2-Mto2 crosslinks identified in our experiments (e.g. between the Mto2 NND and C-terminal region) could be either inter- or intramolecular. However, within our model, either mode of interaction between the Mto2 NND and C-terminal region could stabilize the NND-NND dimer interface (Figure 7; Suppl. Figure 7).

Are there alternatives to the ABBA model that are also consistent with our results? In principle, it is possible that two Mto2 molecules could form a closed bivalent dimer (e.g. as in Suppl. Figure 5B but including additional Mto2-Mto2 interactions identified by CLMS) that multimerizes by binding other closed bivalent Mto2 dimers. However, if this were the case, an explanation of multimerization would require invoking the existence of additional Mto2-Mto2 interaction surfaces (i.e. beyond the NND and the TCD), for which there is currently no experimental evidence. By contrast, the ABBA model provides a parsimonious explanation for Mto2 multimerization. In this context, we note that unambiguously intermolecular Mto2-Mto2 CLMS crosslinks were found essentially only in the NND and the TCD, and in regions that could be crosslinked to them (Figure 5A, 5C). This suggests that these are the key sites of intermolecular Mto2-Mto2 interaction, as proposed in the ABBA model.

Although our experiments used Mto1[bonsai] rather than full-length Mto1 (see Methods), our model for Mto1/2[bonsai] assembly should apply equally well to Mto1/2 complex containing full-length Mto1. Similarly to Mto1[bonsai], MT nucleation by full-length Mto1 *in vivo* is strongly dependent on Mto2 (except at sites like the SPB, where Mto1 is highly concentrated independently of Mto2), and this has been attributed to Mto2 multimerization (Lynch et al., 2014; Samejima et al., 2005; Samejima et al., 2008). Mto1 sequences outside the “bonsai” region mainly include short regions involved in Mto1/2 complex localization (Bao et al., 2018; Samejima et al., 2010) and multiple predicted coiled-coils (Suppl. Figure 1A). Because Mto1[bonsai] already contains several predicted coiled-coils (Suppl. Figure 1A; Figure 5C), it is likely that the coiled-coils of “non-bonsai” regions simply increase the length of the coiled-coil dimer that already exists in Mto1[bonsai]. In any case, even if the additional non-bonsai coiled-coils were to form a slightly higher-order oligomer (e.g. a coiled-coil tetramer), such an oligomer could be further multimerized by binding to multimeric Mto2.

The ABBA model proposes that Mto1/2 complex forms a linear multimer with a range of possible sizes (Figure 7, Suppl. Figure 7). This is consistent with previous observations that GFP-tagged Mto1/2[bonsai] puncta *in vivo* display a range of intensities (**Lynch et al., 2014**). We imagine the Mto1/2 multimer to be open-ended. While it is also possible that the Mto1/2 multimer could close back on itself to form a ring, we consider this unlikely, as *γ*-TuSCs bound to Mto1/2 would be expected to form a helix rather than a ring (see below). Mto1/2 multimer size *in vivo* is likely to be limited by protein abundance and possibly also by internal subunit dissociation, which can serve to regulate the length of simple (i.e. non-multistranded) linear multimers (Howard, 2001). Phosphorylation may be another important regulatory mechanism in the dynamic assembly of Mto1/2 multimers, as we previously showed that Mto2 phosphorylation at multiple sites contributes to Mto1/2 complex instability *in vivo*, in both interphase and mitosis (Borek et al., 2015). Currently we do not know the detailed phosphorylation state of recombinant insect cell-expressed Mto2. Because multiple phosphorylation states of Mto2 coexist throughout the cell cycle *in vivo* ((Borek et al., 2015); see also Figure 6B), it is possible that these different states collectively “tune” aspects of Mto1/2 complex assembly.

In this work we have not addressed how the Mto1/2 complex interacts with the *γ*-TuC. Previously it was shown by cryo-electron microscopy that purified budding yeast *γ*-TuSCs, together with a fragment of the CM1 protein Spc110p, can associate *in vitro* to form a low-pitch helix with near-correct orientations for MT nucleation (Kollman et al., 2015; Kollman et al., 2010). Recently we demonstrated that Mto1/2[bonsai], fission yeast *γ*-TuSC, and the small protein Mzt1 together form active MT-nucleation complexes *in vitro* (Leong et al., 2019). Based on the budding yeast work, we favor the idea that disordered regions of Mto2 could allow the multimeric Mto1/2 complex to serve as a flexible “landing strip” that brings *γ*-TuSCs into proximity and allows them to interact without constraining their intrinsic geometry of interaction. Interestingly, while Mto1/2[bonsai] puncta *in vivo* exhibit a wide range of fluorescence intensities, the puncta that actively nucleate MTs have a smaller range of intensities (Lynch et al., 2014). Therefore, in the context of an open-ended Mto1/2 assembly mechanism, it is possible that Mto1/2 interaction with *γ*-TuSCs could also contribute to regulating Mto1/2 multimerization state.

It is instructive to compare the design principles for Mto1/2 assembly suggested by our work with the principles proposed for CM1 proteins at the centrosome in animal cells, as there may be a link between the requirements for MT organization in a given cell type and the mechanisms used to organize that cell’s MTOCs. For example, the *Drosophila* CM1 protein Cnn, which recruits the *γ*-TuC to the centrosome, forms micron-scale 2-D and/or 3-D assemblies *in vitro* and at the centrosome *in vivo* (Citron et al., 2018; Conduit et al., 2014; Feng et al., 2017). These assemblies are generated via combined homotypic and heterotypic inter- and intramolecular interactions involving non-CM1 domains that are conserved in human cells. Formation of 2-D or 3-D assemblies requires high internal subunit connectivity (i.e. individual subunits interacting with three or more other subunits; (Yeates, 2017)). High connectivity confers additional stability to the assembly, and in an open-ended assembly this in turn can lead to most of the cell’s CM1 proteins accumulating at a single site. In cells in which the centrosome is the single predominant MTOC and nucleates hundreds of MTs, such an assembly makes biological sense, because accumulation of nucleation proteins at the centrosome not only promotes nucleation at that site but also prevents nucleation at non-centrosomal sites. By contrast, in a typical interphase fission yeast cell there are only ∼10-30 MTs in total, organized as independent linear arrays of MT bundles (Sawin and Tran, 2006), and MTs are nucleated by multiple non-centrosomal MTOCs dynamically distributed throughout the cell, with many MTOCs nucleating only a single MT. In this instance, building smaller-scale CM1 protein assemblies with lower internal connectivity (i.e. each subunit interacting with only two other subunits, to form a linear structure) may be more suited to their biological function, allowing the formation and coexistence of multiple independent MTOCs.

## ACKNOWLEDGEMENTS

We thank I. Berger for MultiBac plasmids, M. Wear and M. Nowicki for help with SEC-MALS, L. Blackburn for help in ITC and Thermal Denaturation Assay, D. Kelly for help with fluorescence microscopy, C. Combe and L. Kolbowski for discussions on crosslinking mass spectrometry, and members of our laboratories for discussion. We thank the Edinburgh Protein Production Facility for support with protein purification and biophysical characterization, the University of Edinburgh Xtalpod facility for crystallization screening, and beam-line scientists at Diamond Light Source for support during data collection. We thank A. Leech (Bioscience Technology Facility, University of York) for assistance with some of the SEC-MALS experiments. Structural coordinates for the Mto2 TCD have been deposited in the PDB with the following accession code: 6GDJ. MS crosslinking data have been deposited in the PRIDE database with the dataset identifier PXD015086. This work was supported by the Wellcome Trust ([094517], [210659] to KES, [108504] to JR, [200898] to AGC, Wellcome Trust Multi-User Equipment grant [101527] for the Edinburgh Protein Production Facility), by Wellcome-University of Edinburgh Institutional Strategic Support Fund (ISSF) Awards for SEC-MALS and crystallization facilities, and by the UK Medical Research Council ([G1000520/1] to AGC). EML and XXB were supported by Darwin Trust of Edinburgh studentships. WEB was supported by a Cancer Research UK studentship (C20060/A10789). The Wellcome Centre for Cell Biology is supported by core funding from the Wellcome Trust [203149]

## MATERIALS AND METHODS

### Baculovirus plasmids and bacmids

Recombinant protein expression in insect cells used the MultiBac baculovirus system, with genes initially cloned into the vector pFL (Trowitzsch et al., 2010) (pKS1219 in our collection). Cloning experiments used *E. coli* DH5*α*, and plasmids were confirmed by sequencing. Plasmids and oligonucleotides used are listed in **Suppl. Data File 1**.

To construct a plasmid for 6xHis-tagged Mto1[bonsai] expression, the sequence for 6xHis-TEV-Mto1[bonsai] (containing Mto1 residues 131-549) was first amplified by PCR, using oligonucleotide primer pair OKS2182/OKS2184, and blunt-cloned into pJET1.2 (Thermo Fisher Scientific), to make pKS1217. The 6xHis-TEV-Mto1[bonsai] sequence from pKS1217 was then subcloned into the pFL P10 promoter multiple cloning site (MCS), using NcoI/NsiI sites, to make pKS1222. MultiBac plasmids for expression of 6xHis-tagged Mto1/2[bonsai] (pKS1225), 6xHis-tagged Mto2 (pKS1258) and GST-6xHis-tagged Mto1[bonsai] (pKS1548) were recently described (Leong et al., 2019). We used Mto1[bonsai] rather than full-length Mto1 in our experiments because Mto1[bonsai] is sufficient for Mto1-dependent MT nucleation *in vivo* (Lynch et al., 2014) and *in vitro* (Leong et al., 2019), and preliminary experiments with baculovirus-expressed full-length Mto1 revealed extensive protein degradation.

A plasmid for expression of mutant Mto2-134AAA was generated by PCR using mutant plasmid pKS1235 (see below, under “Fission yeast strain construction”) as template and oligonucleotide primer pair OKS2320/OKS2186. The PCR product was digested with BclI and SalI and cloned into BamHI/SalI sites of pFL (BclI was used because the Mto2 coding sequence contains internal BamHI sites). The resulting plasmid was named pKS1310. A plasmid for expression of mutant Mto2-134A136A was generated by PCR-based site-directed mutagenesis, using pKS1258 as template and oligonucleotide primer pair OKS2823/OKS2824), to make pKS1543.

Plasmids containing internal deletions and N- and C-terminal truncations of Mto2 were generated by inside-out PCR amplification of pKS1258, followed by phosphorylation with T4 polynucleotide kinase and ligation. Oligonucleotide primer pairs OKS3792/OKS3791, OKS3794/OKS3793, OKS3796/OKS3795, OKS3799/OKS3797, and OKS3798/OKS3799 were used to make plasmids pKS1717 (Mto2[44-397]), pKS1718 (Mto2[ΔNND]), pKS1719 (Mto2[ΔTCD]), pKS1720 (Mto2[1-319]), and pKS1721(Mto2[1-350]), respectively.

Recombinant bacmids were generated via Tn7 transposition, by transformation of DH10 EMBacY *E. coli* cells (Trowitzsch et al., 2010) with the pFL-based plasmids described above. Transformed cells were plated on triple-antibiotic plates (50 µg/mL kanamycin, 7 µg/mL gentamycin, 12.5 µg/mL tetracycline) containing 0.2 mM IPTG, and 200 µg/mL 5-bromo-3-indolyl-beta-galactoside (Bluo-Gal) and screened for white colonies, indicating transposition. White colonies were picked and grown overnight at 37°C in 5-10 ml LB with triple-antibiotic selection, with shaking at 200 rpm. Recombinant bacmid DNA was isolated by isopropanol precipitation, air-dried, re-suspended in 50 μL sterile dH_2_O.

### Baculovirus production

Bacmid DNA was transfected into adherent Sf9 cells (Thermo Fisher Scientific) in six-well culture plates (∼1 x 10^6^ cells per well), in 2-2.5 mL Sf900 II serum-free medium (SFM) supplemented with 1x PenStrep from 100x stock (Life Technologies, Thermo Fisher). For transfection, 45 μL of bacmid DNA was mixed with 240 μL fresh Sf900 II SFM/PenStrep, then supplemented with 15 μL of X-tremeGENE 9 HP transfection reagent (Roche) and mixed well. The complete mixture was incubated at 27°C for 30 min and then slowly added to the six-well plate drop by drop, with gentle shaking by hand. After ∼72 h, cells were collected from six-well plates and transferred to a 100 mm tissue culture dish (#353003; BD Biosciences) containing 10^7^ fresh Sf9 cells in 10 mL Sf900 II SFM/PenStrep. Appearance of YFP-fluorescent cells (due to expression of YFP gene in EMBac Y) during the subsequent 48-72 h was used as an indicator of protein expression and release of functional virus into culture medium. Cells were then dislodged into the medium by pipetting up-down and centrifuged (at 1000 x *g* for 5 min) in a 15 mL falcon tube. Supernatant was collected in a fresh sterile falcon tube and stored with 10% FBS as V0 virus stock (protected from light at 4°C). Pelleted cells were also recovered, to confirm expression (e.g. by Western blotting). Larger-volume, later-generation V1 and V2 virus stocks were produced as described previously (Bieniossek et al., 2008) and stored in the same way as V0 virus.

### Baculovirus protein expression and purification

For large-scale recombinant protein production, fresh V2 virus was generated and used to infect Sf9 suspension culture (cell density 1.5-2.5 x 10^6^/mL) in a 1-2 L volume, with a virus:cell culture ratio (v/v) of 1:10-1:20 (Bieniossek and Berger, 2009). When expressed individually, Mto1[bonsai] and Mto2 were expressed as N-terminal 6-His-tagged proteins. For coexpression, the tag was present only on Mto1[bonsai]. After 48-72 h of infection, depending upon cell viability and YFP fluorescence, cells were harvested by centrifugation at 1100 x *g* for 10 min, washed by gentle resuspension in PBS, and stored as cell pellet at −80°C or used immediately for further purification. Protein expression at each generation of virus amplification was monitored by SDS-PAGE/immunoblotting and/or by YFP fluorescence microscopy.

For purification of Mto2, virus-infected Sf9 cells were resuspended and lysed using a Dounce homogenizer in 50 mM Tris-HCl, pH 8.0, 200 mM NaCl, 5 % glycerol, 5 mM *β*-mercaptoethanol, 45 mM imidazole, 0.1% Triton-X-100, 10 U/mL Base Muncher (Expedeon), 1 mM phenylmethane sulfonyl fluoride (PMSF) and 1x CLAAPE protease inhibitors, from 2000x stock (2000x CLAAPE = 10 mg/mL chymostatin, 10 mg/mL leupeptin, 10 mg/mL antipain, 10 mg/mL aprotinin, 10 mg/mL pepstatin, and 10 mg/mL E-64, in DMSO), using 3 mL buffer per 1 g of cells. Following further lysis by sonication (Soniprep 150, 3 mm probe size, 3 pulses, 30 sec each, at amplitude 15, with 1 min rest interval), the sample was cleared by centrifugation at 25,000 rpm (JA25.50 rotor, Beckman Coulter) for 1.5 h at 4°C. Mto2 was bound in batch using nickel-charged Fractogel EMD chelate(M) (Merck) overnight at 4°C. The resin was packed into a column and washed in 20-50 column volumes (CV) of IMAC-Tris Buffer (50 mM Tris-HCl, pH 8.0, 300 mM NaCl, 5% glycerol (v/v), 5 mM *β*-mercaptoethanol) supplemented with 100 mM KCl, 90 mM imidazole, 10 mM MgCl_2_, and 10 mM ATP. Further wash steps (5 CV) were carried out with IMAC-Tris Buffer supplemented with 100 mM KCl and 90 mM imidazole. Protein was eluted with a series of step elutions of IMAC-Tris Buffer containing increasing imidazole concentrations (100-600 mM Imidazole). Mto2 eluted at 600 mM Imidazole. Further purification was carried out using size exclusion chromatography with a Superose 6 column (GE Healthcare), using IMAC-Tris Buffer but without *β*-mercaptoethanol.

For purification of Mto1[bonsai] and Mto1/2[bonsai] complex, a similar protocol was used, but Tris-HCl was replaced with phosphate (1.5 mM KH_2_PO_4_, 5.1 mM Na_2_HPO_4_, pH 8.0) in all buffers, e.g. IMAC-Phosphate Buffer is identical to IMAC-Tris Buffer apart from containing phosphate instead of Tris-HCl. Mto1[bonsai] and Mto1/2[bonsai] complex both eluted at 300 mM imidazole.

### Copurification of Mto2 mutants with GST-Mto1[bonsai]

For coexpression of Mto2 and Mto2 variants with GST-6xHis-TEV-Mto1[bonsai] (here referred to as “GST-Mto1[bonsai]”), V2 virus stocks were mixed in 4:1 ratio of Mto2:GST-Mto1[bonsai], to ensure overexpression of Mto2 (and variants) relative to GST-Mto1[bonsai]. Sf9 cells were then infected at a (total) virus:cell culture ratio (v/v) of 1:15-1:20 and processed as above.

Cells were lysed by sonication on ice as described above, in GST-Tris300 Buffer (50 mM Tris-HCl, pH 8.0, 300 mM NaCl, 2 mM EDTA, 3 mM DTT) supplemented with 100 mM KCl, 0.1% Triton X-100, 10 U/mL Base Muncher, 2 mM PMSF, and 1x CLAAPE protease inhibitors, using 3 mL buffer per 1 g of cells. Cell lysate (∼50 mL) was cleared by centrifugation at 25,000 rpm for 1 h at 4°C (JA25.50 rotor, Beckman Coulter). An aliquot of clarified lysate was kept separately as an “input” sample. GST-Mto1[bonsai] was captured from clarified lysate by passing twice through a glutathione-agarose column (Sigma, G4510) pre-equilibrated in GST-Tris300 Buffer. The column was then washed with 5 CV of GST-Tris300 Buffer supplemented with 20 mM MgCl_2_ and 10 mM ATP, and proteins were then eluted with GST-Tris300 Buffer supplemented with 20 mM glutathione.

To assay copurification of Mto2 mutants with GST-Mto1[bonsai], SDS-PAGE was used for qualitative assessment, and western blotting for quantitative assessment; SDS-PAGE alone was not sufficient because one mutant, Mto2[1-319], comigrated with a non-specific band in the purification. For SDS-PAGE of clarified cell lysate, equal volumes of lysate were loaded (3 µL for Coomassie Blue staining; 0.1 µL for anti-Mto2 Western blot). Because infection efficiency (and thus protein expression) varied among the different infections, for SDS-PAGE of partially purified GST-Mto1[bonsai] and copurifying Mto2, equal amounts of total purified protein were loaded (15 µg for Coomassie Blue staining; 110 ng for anti-Mto2 western blot). Low loadings for anti-Mto2 western blotting ensured that blot signal for quantification remained in a linear range. For western blots, proteins were transferred to nitrocellulose. After blocking, blots were incubated overnight at 4°C with blot-affinity-purified anti-Mto2 primary antibody (1:50 dilution in TBS plus 4% nonfat dry milk), then with unlabelled GT-34 mouse monoclonal anti-goat secondary antibody (Sigma, G2904), at 1:10,000 dilution, to amplify the signal, and then with IRDye680CW donkey anti-mouse tertiary antibody (LI-COR) at 1:5000 dilution. Blots were imaged using an Odyssey CLx fluorescence imager (LI-COR). For quantification, Mto2 signals on western blots were background-subtracted and quantified using Image Studio Lite (version 5.2.5; LI-COR). The resulting values were then normalized to the cognate GST-Mto1[bonsai] signals in the corresponding Coomassie Blue-stained lanes, and then scaled to arbitrary units in which the amount of wild-type Mto2 copurifying with GST-Mto1[bonsai] has a value of 100. The copurification experiment was performed twice; the first time used Coomassie Blue staining only, while the second time (shown) used Coomassie Blue staining and western blotting.

### E. coli expression plasmids

Cloning experiments used *E. coli* DH5*α*, and plasmids were confirmed by sequencing. Plasmids and oligonucleotides used are listed in **Suppl. Data File 1**.

The Mto2 Near N-terminal Domain (NND; Mto2 residues 78-153) and Twin-Cysteine Domain (TCD; Mto2 residues 180-265) were expressed as GST fusions, with the GST sequence N-terminal to the domain of interest. Plasmids were constructed using Gateway cloning. First, NND and TCD sequences codon-optimized for *E. coli* expression were synthesized by GeneArt (Thermo Fisher Scientific), as plasmids pKS1312 and pKS1313, respectively. The NND and TCD sequences were then subcloned into Gateway donor vector pDONR221 (Thermo Fisher Scientific) through BP recombination (BP clonase II enzyme mix; Thermo Fisher Scientific), to make plasmids pKS1315 and pKS1317. Finally, NND and TCD sequences from these plasmids were subcloned into Gateway expression vector pGGWA (Busso et al., 2005) through LR recombination (LR clonase II enzyme mix; Thermo Fisher Scientific), to make plasmids pKS1318 and pKS1320.

Plasmids for expression of NND mutants NND-134AAA, NND-134A136A, NND-N123C, and NND-R147C were generated by PCR, using the QuickChange Lightning Site-Directed Mutagenesis Kit (Agilent) and pKS1318 as template. Mutant oligonucleotide primer pairs were OKS2611/OKS2612 (NND-134AAA), OKS2821/OKS2822 (NND-134A136A), OKS3163/OKS3164 (NND-N123C) and OKS3165/3166 (NND-R147C). The resulting plasmids were named pKS1340, pKS1528, pKS1535, and pKS1536, respectively.

### E. coli protein expression and purification

Proteins were expressed in BL21(DE3) pRIL cells (Agilent), grown in 4 L of LB medium with 100 µg/mL ampicillin and 25 µg/mL chloramphenicol. Cells were grown at 37°C with shaking at 200 rpm to an optical density (600 nm) of 0.7-0.8, cooled to 20°C and induced with 0.2 mM isopropyl β-D-1-thiogalactopyranoside at 20°C overnight. Cells were harvested by centrifugation at 7000 rpm (4°C; 10 min), washed once with PBS, and recentrifuged. The cell pellet was stored frozen at −80°C before lysis and purification. Pellets were resuspended in GST-Tris200 Buffer (50 mM Tris-HCl pH 8.0, 200 mM NaCl, 2 mM EDTA, 3 mM DTT), supplemented with 50 μg/mL lysozyme, 10 U/mL Base Muncher, 2 mM PMSF, and 1x CLAAPE protease inhibitors. Cells were lysed by sonication and the lysate cleared by centrifugation (25,000 rpm, 4°C, 1.5 h, JA 25.50 rotor). The soluble fraction was applied to a 20 mL glutathione-sepharose column pre-equilibrated in GST-Tris200 Buffer and washed with 5 CV of GST-Tris200 Buffer supplemented with 10 mM MgCl_2_ and 10 mM ATP. Proteins were eluted with GST-Tris200 Buffer supplemented with 20 mM reduced glutathione. Eluted proteins were treated with TEV protease to remove the GST tag and separated by size-exclusion chromatography using GST-Tris200 Buffer. Pooled fractions of Mto2 fragments were further separated from GST by a second pass over glutathione-sepharose.

To generate selenomethionine-labeled protein, Mto2 TCD was expressed in B834(DE3) cells in 2xM9 medium supplemented with all amino acids except methionine, which was replaced by selenomethionine (Ramakrishnan et al., 1993). Subsequent purification steps were as above.

SDS-PAGE of NND-N123C and NND-R147C under reducing and non-reducing conditions was performed once.

### Analytical SEC and SEC-MALS

Size-exclusion chromatography (SEC) was performed at 4°C, using a Superose 6 10/300 column (GE Healthcare) for full-length oligomeric proteins and a Superdex 75 10/300 column (GE Healthcare)for Mto2 NND and TCD fragments. Columns were equilibrated with IMAC-Tris Buffer (without *β*-mercaptoethanol), IMAC-Phosphate Buffer (without *β*-mercaptoethanol) or GST-Tris200 Buffer, depending on the protein being analyzed (see above). Columns were loaded with 100 µL of purified protein. Peak elution fractions were analysed on SDS-PAGE with Coomassie Blue staining or by western blotting. Coomassie Blue-stained gels in Fig. 1C,D were scanned in grayscale and later pseudocolored.

Where appropriate, western blots of eluted SEC fractions were probed with homemade sheep anti-Mto1 antiserum (Sawin et al., 2004) or sheep anti-Mto2 antiserum (Samejima et al., 2005), both at 1:1000 dilution. Blots were further incubated with unlabelled GT-34 mouse monoclonal anti-goat antibody, at 1:10,000 dilution, and blots were then probed with IRDye680CW donkey anti-mouse antibody (LI-COR) at 1:5000 dilution. Blots were imaged using an Odyssey fluorescence imager (LI-COR) and analyzed with Image Studio (LI-COR).

For size-exclusion chromatography with multi-angle light scattering (SEC-MALS) experiments, purified proteins were analysed on either a Superose 6 10/300 column (for full-length proteins and protein complexes) (GE Healthcare) or a Superdex 75 10/300 column (for protein fragments) (GE Healthcare) coupled to using a Viscotek SEC-MALS 20 and refractive index detector VE3580 (Malvern Instruments Ltd.). Data were analysed using OmniSEC software (Malvern Instruments Ltd.). Bovine serum albumin (2 mg/mL) was used for normalization of the light-scattering detectors and alignment of UV_280_ and light scattering signal with differential refractive index signal.

For some SEC-MALS experiments (carried out at Bioscience Technology Facility, University of York), proteins were analyzed using a Wyatt HELEOS-II multi-angle light scattering detector and a Wyatt rEX refractive index detector, linked to a Shimadzu HPLC system (SPD-20A UV detector, LC20-AD isocratic pump system, DGU-20A3 degasser and SIL-20A autosampler). Data were analysed using Astra V software (Wyatt).

Data from the same experiment were used for NND SEC-MALS plots shown in Figure 2A and at “600 µM” in Suppl. Figure 2D. This is not obvious because the units on the axes in the two figures are different (because of the need to use MALS 90° signal and derived retention volume in Suppl. Figure 2D). Similarly, data from the same experiment were used for TCD SEC-MALS plots in Fig. 2D and at “1000 µM” in Suppl. Figure 2E.

SEC-dilution experiments were performed twice for all proteins and fragments. SEC-MALS experiments were performed twice for Mto1/2[bonsai] complex, Mto1[bonsai], Mto2, Mto2-134A136A, and Mto2-134AAA, once for NND-134A136A and NND-134AAA, and once for each dilution series of the NND and the TCD. SEC after mixing Mto1[bonsai] with Mto2 was performed twice. For experiments with multiple replicates, representative examples are shown.

### Thermal denaturation assay

Purified Mto2 TCD (45 µL of 20 µM purified protein in GST-Tris200 Buffer) was mixed with 5 µL of 1x SYPRO Orange dye (Thermo Fisher Scientific) and subjected to a temperature (20-95°C) vs fluorescence scan on an iQ5 Real-Time PCR system (BioRad). Increased fluorescence represents SYPRO Orange dye binding to hydrophobic regions of the TCD that become exposed upon temperature-dependent unfolding. The experiment was performed once, taking the average value from triplicate samples.

### Quantification of endogenous Mto2 expression

For quantification of endogenous fission yeast Mto2 expression, wild-type and *mto2Δ* cells were grown (separately) to a density of ∼1.2×10^7^ cells/mL, harvested by centrifugation at 4000 rpm for 4 min at room temperature, washed in TBS plus 2 mM PMSF and 2 mM EDTA, and heated at 95°C for 5 min to denature proteases (Bicho et al., 2010). Cells were then lysed by bead-beating with 0.5 mm zirconia beads (Biospec, #11079105z) in a Ribolyser (Hybaid) for 2×30 s at speed “6.5”. The supernatant was collected, mixed with an equal volume of 2X Laemmli sample buffer without bromophenol blue or reducing agent, and heated again at 95°C, followed by centrifugation for 10 min at 13,000 rpm and recovery of the supernatant. Protein concentration was determined by BCA assay (Thermo Fisher Scientific), and lysates were then supplemented with DTT (0.1 M final) and bromophenol blue (0.01% final). Next, known amounts of recombinant purified 6xHis-Mto2 (0.25, 0.5, 1.0, 2.0, 3.0, 4.0 ng) were added to a volume of *mto2Δ* cell lysate corresponding to 50 µg total protein, and samples were separated by SDS-PAGE alongside a volume of wild-type cell lysate also corresponding to 50 µg total protein. Proteins were transferred to nitrocellulose and incubated with sheep anti-Mto2 antiserum (1:1000), processed as described above for SEC fractions, and analyzed quantitatively using Image Studio (LI-COR). Signals from *mto2Δ* lysates containing different amounts of recombinant 6xHis-Mto2 protein were used to generate a standard curve of western blot signal as a function of the amount of Mto2, and this was then used to calculate the amount of Mto2 present in the wild-type cell lysate. The determined value of ∼2.3 ng Mto2 per 50 µg wild-type cell lysate corresponds to ∼2 x 10^-16^ g Mto2 per cell. Based on average cell volume and the proportion of cell volume occupied by endomembrane compartments (29%; (Wu and Pollard, 2005)), this is equivalent to a cytosolic Mto2 concentration of ∼160 nM. The western blotting was performed twice, using the same samples each time.

### Chemical crosslinking & LC-MS/MS (CLMS)

Crosslinking experiments were carried out in a buffer containing 3 mM KH_2_PO_4_, 10.2 mM Na_2_HPO_4_, pH7.5, 300 mM NaCl, 5% glycerol, using the zero-length crosslinker, 1-ethyl-3-(3-dimethylaminopropyl) carbodiimide hydrochloride (EDC, Thermo Fisher Scientific) in the presence of N-hydroxysulfosuccinimide (Sulfo-NHS, Thermo Fisher Scientific). First, proteins were dialyzed against a buffer containing 1.5 mM KH_2_PO_4_, 5.1 mM Na_2_HPO_4_, pH7.5, 300 mM NaCl, 5% Glycerol, and then an appropriate volume of buffer with a 20-fold higher phosphate concentration (30 mM KH_2_PO4, 102 mM Na_2_HPO_4_, pH7.5, 300 mM NaCl, 5% glycerol) was added achieve the desired phosphate concentration. Proteins were then mixed with crosslinker (20-25 µg Mto2 with 22 µg EDC and 44 µg Sulfo-NHS, or 30 µg of Mto1/2[bonsai] complex with 10 µg EDC and 20 µg Sulfo-NHS, both in a total volume of 30 µL, or 6 µg Mto1[bonsai] with 10 µg EDC and 20 µg Sulfo-NHS, in a total volume of 20 µL) and incubated for 2 h at 18°C (Mto2 and Mto1/2[bonsai] complex) or 1 h at 18°C (Mto1[bonsai]).

Crosslinked protein bands were resolved on NuPAGE Tris-Acetate gels (Thermo Fisher Scientific) and visualised with Coomassie Blue stain. Gel bands corresponding to monomer and higher oligomers of crosslinked complexes were excised, in-gel reduced and alkylated, then digested by trypsin following a standard protocol (Maiolica et al., 2007). The peptide mixture was then desalted using C18-Stage-Tips (Rappsilber et al., 2003) for LC–MS/MS analysis.

Samples were analysed using an UltiMate 3000 Nano LC system coupled to an Orbitrap Fusion Lumos Tribrid mass spectrometer (Thermo Scientific), applying a ‘high-high’ acquisition strategy. Peptides were separated on a 50 cm C18 (75 μm ID, 2 μm particles, 100 Å pore size) PepMap EASY-Spray column (Thermo Scientific) fitted into an EASY-Spray source (Thermo Scientific), operated at 55 °C column temperature. Mobile phase A consisted of water and 0.1% v/v formic acid. Mobile phase B consisted of 80% v/v acetonitrile and 0.1% v/v formic acid. Peptides were loaded at a flow rate of 0.3 µL/min and eluted at 0.2 μL/min using a linear gradient going from 2% mobile phase B to 40% mobile phase B over 109 min, followed by a linear increase from 40% to 95% mobile phase B in 11 min. MS data were acquired in the data-dependent mode with a scan range of 300-1700 m/z. For each 3 s acquisition cycle, the survey level spectrum was recorded in the Orbitrap with a resolution of 120,000. The ions with a precursor charge state between 3-8 and an intensity higher than 5×10^4^ were isolated and fragmented using high energy collision dissociation (HCD) with collision energy 30% according to charge order. The fragmentation spectra were recorded in the Orbitrap with a resolution of 15,000. Dynamic exclusion was enabled with single repeat count and 60 s exclusion duration.

The mass spectrometric raw data were processed to generate peak lists by MSCovert (ProteoWizard 3.0.11417) (Kessner et al., 2008), and crosslinked peptides were matched to spectra using Xi software (version 1.6.731) (Mendes et al., 2019) with in-search assignment of monoisotopic peaks (Lenz et al., 2018). The search parameters were as follows: MS accuracy, 3 p.p.m.; MS2 accuracy, 10 p.p.m.; enzyme, trypsin; allowed number of missed cleavages, 4; crosslinker, EDC; missing mono-isotopic peaks, 2; fixed modifications, carbamidomethylation on cysteine; variable modifications, oxidation on methionine, phosphorylation on serine, threonine; fragments, b and y ions with loss of H_2_O, NH_3_ and CH_3_SOH. FDR was computed using XiFDR and results reported at 5% residue level FDR (Fischer and Rappsilber, 2017).

Mto2 CLMS experiments were performed twice, with one experiment generating one MS sample and the other generating two MS samples. Raw data from the three MS runs were pooled for analysis. Mto1/2[bonsai] CLMS experiments were performed three times, with each experiment generating two MS samples. Raw data from the six MS runs were pooled for analysis.

The mass spectrometry proteomics data have been deposited to the ProteomeXchange Consortium via the PRIDE (Perez-Riverol et al., 2019) partner repository with the dataset identifier PXD015086.

### Isothermal titration calorimetry (ITC)

Purified Mto2 NND was buffer-exchanged into ITC Sample Buffer (50 mM Tris-HCl pH 8.0, 150 mM NaCl, 2 mM EDTA) by size exclusion chromatography. ITC dilution-series experiments were carried out at 25°C using an iTC200 system (Microcal; Malvern Instruments). Heat change (endothermic pulse) upon dilution was measured by stepwise injection (2 µL each, 20 injections) of protein sample (100 µM, 40 µL volume) into cell containing ITC Sample Buffer (200 µL volume). The resulting heat changes were integrated using Origin software (OriginLab) and curve fitting was performed to determine the dissociation constant and enthalpy (ΔH) using two-steps dimer dissociation model provided by the software package (McPhail and Cooper, 1997). The experiment was performed three times; a representative example is shown.

### Protein crystallization and structure determination

Mto2 TCD crystals were grown using the vapor-diffusion method. Crystals of native protein (45 mg/mL) were grown in 20% PEG 3350, supplemented with 0.2 M sodium thiocyanate, pH 7. Crystals of selenomethionine-labeled protein (17 mg/mL) were grown in 20 % PEG 3350, with 0.2 M NH_4_Cl, pH 6.6, using streak-seeding with native crystals. Native crystals were hexagonal (*P*6_1_) while selenomethionine-labeled crystals were orthorhombic (*P*2_1_2_1_2_1_). Crystals were cryoprotected in their corresponding crystallization buffers supplemented with 30 % glycerol and flash cooled to 100° K. Selenomethionine diffraction data were indexed, integrated and scaled using the xia2 package (Winter, 2010; Winter et al., 2013) and phases were determined using SHELX (Sheldrick, 2008). Model building was carried out in COOT (Emsley et al., 2010) with refinement in REFMAC (Murshudov et al., 2011; Winn et al., 2011). The refined model was then used as a search model for the native dataset in PHASER (McCoy et al., 2007; Winn et al., 2011) with subsequent refinement in REFMAC. The final model includes 71 residues out of a total of 86 residues that were present in the crystal. The C-terminal 17 residues could not be modelled due to poor electron density in these regions that made a clear interpretation of the map difficult. Geometrical parameters were checked using MOLPROBITY (Chen et al., 2010), and structural data were subsequently analyzed using PDBePISA (Krissinel and Henrick, 2007) and PDBeFOLD (Krissinel and Henrick, 2004).

### Fission yeast strain construction

Fission yeast strains were constructed by targeted plasmid integration (Fennessy et al., 2014), PCR-based gene targeting (Bahler et al., 1998), and genetic crosses (Ekwall and Thon, 2017). Standard methods for were used culture (Petersen and Russell, 2016). Cells were grown either in YE5S rich medium, using Bacto yeast extract (Becton Dickinson) or in EMM2 or EMMG defined minimal medium. Supplements for auxotrophies were used at 175 mg/L. Antibiotics (G418, nourseothricin, and hygromycin) were used at 100 µg/mL. Strains used in this work, and plasmids and oligonucleotides used in strain construction, are listed in **Suppl. Data File 1**.

Mo2 NND and TCD domains were overexpressed (i.e. in separate strains) from transgenes integrated at the *hph.171k* locus on chromosome III by homologous recombination (Fennessy et al., 2014). NND and TCD sequences were amplified by PCR using plasmid pKS1258 as template and oligonucleotide primer pairs OKS3315/OKS3316 (for NND) and OKS3313/OKS3314 (for TCD). PCR products were cloned into pJET1.2 (Thermo Fisher Scientific) by blunt-end ligation, to generate plasmids pKS1572 and pKS1573. NND and TCD inserts from these plasmids were then subcloned into fission yeast integration plasmid pINTH1 (Fennessy et al., 2014) (pKS1612 in our collection), using NdeI and BamHI sites, to generate plasmids pKS1574 and pKS1579.

The pINTH-based NND and TCD plasmids pKS1574 and pKS1579 were linearized using NotI and transformed (separately) into yeast strain KS7742. Colonies that had integrated plasmids at the *hph.171k* locus (leading to interruption of the *hph* coding sequence) were identified by simultaneous acquisition of nourseothricin resistance and loss of hygromycin resistance. Strains were further confirmed by yeast colony PCR, using oligonucleotide primer pairs OKS253/OKS3174 for the NND and OKS253/OKS3172 for the TCD. The resulting strains were named KS8641 (for NND overexpression) and KS8642 (for TCD overexpression). On minimal-medium agar plates (i.e. under derepressing conditions, in absence of thiamine), KS8641 and KS8642 showed a curved cell phenotype, consistent with impaired Mto2 function.

To introduce *mto1Δ* and *mto2-GFP* into strains KS8641 and KS8642, these strains were crossed successively with strains KS7055 and KS9111, to generate strains KS9196 (for NND overexpression) and KS9200 (for TCD overexpression).

To generate *nmt81:*Mto2 C-terminal truncations tagged with mECitrine, we first constructed the strain KS9608, in which the *mto2* promoter at the endogenous *mto2* locus was replaced by the *nmt81* promoter and an upstream *natMX6* antibiotic resistance marker. KS9608 was constructed by replacing the *kanMX6* resistance marker in strain KS1474 *(kanMX6:nmt81:mto2)* with the *kanMX6* resistance marker, via transformation with a PCR containing the *natMX6* cassette, followed by homologous recombination. Next, KS9608 was transformed with different PCR products, to generate full-length *nmt81:*Mto2 tagged at its C-terminus with mECitrine, or *nmt81:*Mto2 C-terminal truncations tagged at their C-termini with mECitrine. For the PCR, plasmid pFA6a-mEcitrine-kanMX6 (Ye et al., 2012) (pKS1330 in our collection) was used as template, with oligonucleotide forward primers OKS257 (full-length Mto2), OKS3908 (Mto2[1-319]), OKS3909 (Mto2[1-340]), OKS3910 (Mto2[1-349]), and OKS3911 (Mto2[1-374]), and oligonucleotide reverse primer OKS3919 for all truncations. After transformation, colonies were selected for simultaneous nourseothricin and G418 resistance (for *nmt81* promoter and mECitrine tag, respectively) and confirmed by microscopy. The resulting strains were named KS9632, KS9621, KS9622, KS9623, and KS9624, respectively (i.e. as in order of different forward primers above). To introduce *mto1Δ* into these strains, they were then crossed with strain KS7353. Colonies from tetrads were screened for nourseothricin and G418 resistance, and fluorescent-protein fusions were confirmed by microscopy. *mto1Δ* was confirmed by colony PCR, using oligonucleotide primer pairs OKS3604/OKS504. The resulting strains were named KS9649, KS9643, KS9644, KS9646, and KS9648, respectively.

To generate *nmt81:*Mto2 N-terminal truncations tagged with mECitrine, the C-terminus of the wild-type *mto2* coding sequence at the endogenous mto2 locus was first tagged with mECitrine by PCR-based gene targeting into strain KS515. PCR used plasmid pKS1330 as template and oligonucleotide primers OKS257/OKS258, using a PCR fragment and homologous recombination, to make strain KS9168. Next, KS9168 was transformed with different PCR products containing the *nmt81* promoter plus homology to different N-terminal sequences, to generate a series of mECitrine-tagged *nmt81:*Mto2 N-terminal truncations by homologous recombination. For the PCR, plasmid pFA6a-natMX6-P81nmt1 (Van Driessche et al., 2005) (pKS706 in our collection) was used as template. Forward primer for all truncations was OKS2413, and reverse primers were OKS4047 (Mto2[24-397), OKS4048 (Mto2[44-397), OKS4049 (Mto2[59-397), OKS4050 (Mto2[69-397), OKS4051 (Mto2[78-397), OKS4052 (for Mto2[90-397). Correct integration of PCR products was confirmed by colony PCR using oligonucleotide primers OKS3924/OKS1804. The resulting strains were then crossed with strain KS7352, to introduce *mto1Δ*. Genotypes from crosses were confirmed by colony PCR, using primer pairs OKS3604/OKS504 for *mto1Δ* and OKS4065/OKS4068 for *nmt81*:Mto2 N-terminal truncations. The resulting truncation strains were named KS9968, KS9969, KS9971, KS10078, KS10009, and KS9973 respectively (i.e. as in order of different reverse primers above).

To construct site-directed mutations in the *mto2* NND and TCD, a *hphMX6* hygromycin resistance marker was first integrated in the *mto2* 3’ untranslated region by transformation of wild-type strain KS516 with a PCR product and homologous recombination. PCR used template plasmid pFA6a-hphMX6 (Hentges et al., 2005) (pKS694 in our collection) and oligonucleotide primer pair OKS2304/OKS2305. The integration site was confirmed by colony PCR using primer pairs OKS1355/OKS2326 and OKS1355/OKS2327. The resulting strain was named KS6901. The presence of the *hph:MX6* marker adjacent to the *mto2* coding sequence did not affect Mto2 protein expression or function (checked by western blot and cell shape, respectively) and was used to help identify *mto2* mutants in subsequent steps. Site-directed mutants were made by a three-step process. In the first step, the *mto2* coding sequence in KS9601 was deleted and replaced by the *ura4+* gene, using PCR and homologous recombination. PCR used template plasmid KS-ura4 (Bahler et al., 1998) (pKS131 in our collection) and oligonucleotide primer pair OKS2302/OKS2303. Colonies were selected for stable uracil prototrophy, and *mto2Δ* was confirmed by western blotting. The resulting strain was named KS6924. In the second step, Mto1[bonsai]-GFP and mCherry-Atb2 were introduced by crossing strain KS6924 with strain KS6678, to make strain KS6928. In the third step, KS6928 was transformed with different PCR products to replace the *ura4+* gene at the *mto2* locus with full-length mutant *mto2* sequences, by homologous recombination. For the PCR, template plasmids pKS1229 (*mto2-103AAA*), pKS1230 (*mto2-112AA116AA*), pKS1234 (*mto2-129AAA*), pKS1231 (*mto2-129AAA134AAA*), pKS1235 (*mto2-134AAA*), and pKS1238 (mto1-103AAA) were synthesized by GeneArt and amplified using oligonucleotide primer pair OKS2344/OKS2345. After transformation, cells were grown on plates containing 0.2% 5-fluoroorotic acid, to select for replacement of *ura4+* by mutant *mto2*. The final strains were named KS7061, KS7063, KS7071, KS7065, KS7073, and KS7079, respectively (i.e. as in order of different template plasmids above).

### Microscopy

Microscopy experiments used a spinning-disk confocal microscope consisting of a TE2000 microscope base with 100x/1.45 NA Plan Apo objective (Nikon), a modified Yokogawa CSU-10 spinning disk module (Visitech), an Optospin IV filter wheel (Cairn Research) and an iXon+ Du888 EMCCD camera (Andor). Stage position and focus were controlled by a MS-2000 automated stage and a CRISP autofocus system (ASI). Temperature was maintained at 25°C using a thermo-regulated chamber (OKOlab). Illumination and data acquisition were controlled by Metamorph software (Molecular Devices).

All imaging experiments were performed in EMMG minimal medium with appropriate supplements. Cell cultures were grown to mid-log phase at 25°C with or without 15 µM thiamine as needed; for experiments involving derepression by removal of thiamine, cells were grown in medium without thiamine for 44-48 h prior to imaging. Imaging was performed using cells mounted on medium-agarose pads, as described (Snaith et al., 2010).

GFP-fusion proteins were illuminated with 488 nm laser light, and mECitrine-fusion proteins with 514 nm laser light. Images were acquired as stacks of 8-11 Z-sections, with 0.6 µm spacing and 0.13 x 0.13 µm pixel size. Image processing was done using ImageJ and Photoshop (Adobe). Images shown in figures are maximum projections of Z-stacks.

Imaging experiments were performed once for each strain/condition (except for wild-type controls, which were included in multiple imaging sets), and multiple images were acquired for each strain/condition Representative images are shown. For quantification of over-curved MTs and number of MT bundles per cell, 50 cells were scored for each strain.

### Fission yeast coimmunoprecipitation

For experiments assaying coimmunoprecipitation of Mto2 (wild-type and mutant) and Gtb1 with Mto1[bonsai]-GFP in *mto2* mutants (Figure 6), yeast cells were grown to a density of ∼2×10^7^ cell/mL and harvested by centrifugation at 5000 rpm for 8 min at 4°C. The cell pellet was washed with 10 mM sodium phosphate pH 7.5, 0.5 mM EDTA, recentrifuged, weighed, and resuspended in 10 mM sodium phosphate pH 7.5, 0.5 mM EDTA, using 0.3 mL buffer per 1 g cell pellet. The cell suspension was then frozen drop-wise into liquid nitrogen and stored at −80°C. Frozen cells were subjected to cryo-grinding in a Retsch RM100 electric mortar-grinder [Retsch, Germany] that was precooled with liquid nitrogen. Frozen cell powder was stored at −80°C.

For each IP (i.e. for each strain), 2 g frozen cell powder was resuspended in 3 mL IP buffer (25 mM sodium phosphate, pH 8.0, 100 mM KCl, 0.5 mM EDTA, 0.2% TX-100, 10 μg/ml CLAAPE protease inhibitors, 2 mM AEBSF, 2 mM benzamidine, 2 mM PMSF, 50 mM Na *β*-glycerophosphate, 1 mM NaF, 0.1 mM sodium orthovanadate, 50 nM calyculin A, 50 nM okadaic acid). Cell lysates were clarified by centrifugation at 4000 rpm, once for 5 min and then twice for 15 min. Next, 50 µl of clarified cell lysate was set aside as “input”, and the remainder was mixed with ∼50 µL of Protein G Dynabeads (Thermo Fisher Scientific) that had been pre-coupled to anti-GFP antibody (Borek et al., 2015). The bead-lysate mixture was then incubated with rotation for 1.5 h at 4°C. Cell lysate was then discarded, and beads were washed once with 3 mL IP buffer and once with 1 mL IP buffer, transferred to a microfuge tube, and washed twice with 0.5 mL IP buffer. Beads were then resuspended in 40 µL Laemmli sample buffer without bromophenol blue or reducing agent and eluted by incubation at 50°C for 15 min, vortexing halfway through the incubation. Eluates from beads were transferred to a fresh microfuge tube, supplemented with DTT (0.1 M final) and bromophenol blue (0.01% final), and heated at 95°C for 5 min. For SDS-PAGE, gels were loaded with 8 µL of clarified cell lysate (input), and 8-12 µL of IPs, depending on the protein to be analyzed by western blot.

For western blots, proteins were transferred to nitrocellulose in 10 mM CAPS (pH 11) and 10% methanol. Equal loading was confirmed by Ponceau S stain. Western blots were blocked with TBS containing 0.025% Tween 20 plus 2% nonfat dry milk and incubated overnight with sheep anti-Mto1 antiserum (1:1000), sheep anti-Mto2 antiserum (1:1000), or mouse monoclonal anti-*γ*-tubulin antibody GTU-88 (1:4000 for cell lysate, 1:1000 for IP; Sigma, T6557), all in blocking solution. For blots involving sheep primary antibodies, blots were incubated with mouse anti-sheep/goat monoclonal antibody GT-34 (1:30,000) for 1 h. All blots were then incubated with IRDye800CW-labeled donkey anti-mouse antibody (1:10,000; LI-COR, 926-32212) for 1 h. After washing in TBS/Tween and then in TBS, blots were imaged on an Odyssey fluorescence imager (LI-COR) and quantified using Image Studio (LI-COR) and Excel (Microsoft) software. Quantification was done slightly differently for Mto2 vs. Gtb1. For Gtb1, the IP lane from cells expressing untagged, full-length Mto1 (left-most lane in anti-GFP IP lanes in Figure 6B) was used as a background lane. For Mto2, the IP lane from *mto2Δ* cells contained a weak, non-specific band that comigrated with Mto2 but did not appear to be due to spillover from an adjacent lane. Because this band was not present in the IP lane from cells expressing untagged, full-length Mto1, it was not appropriate to use the latter as a background lane; therefore, the IP lane from *mto2Δ* cells was used as a background lane. For all strains, the IP signal from the appropriate background lane was subtracted from the IP signal for a given strain, to give a corrected IP signal. This was then normalized to the Mto1[bonsai]-GFP IP signal for the same strain, to give a normalized corrected IP signal. In Figure 6, values for normalized corrected IP signals are presented as a percentage of the normalized corrected IP signal for wild-type cells (set to 100%). Coimmunoprecipitation experiments and accompanying western blots were performed once for each strain.

## SUPPLEMENTARY FIGURE LEGENDS

**Supplementary Figure 1.**
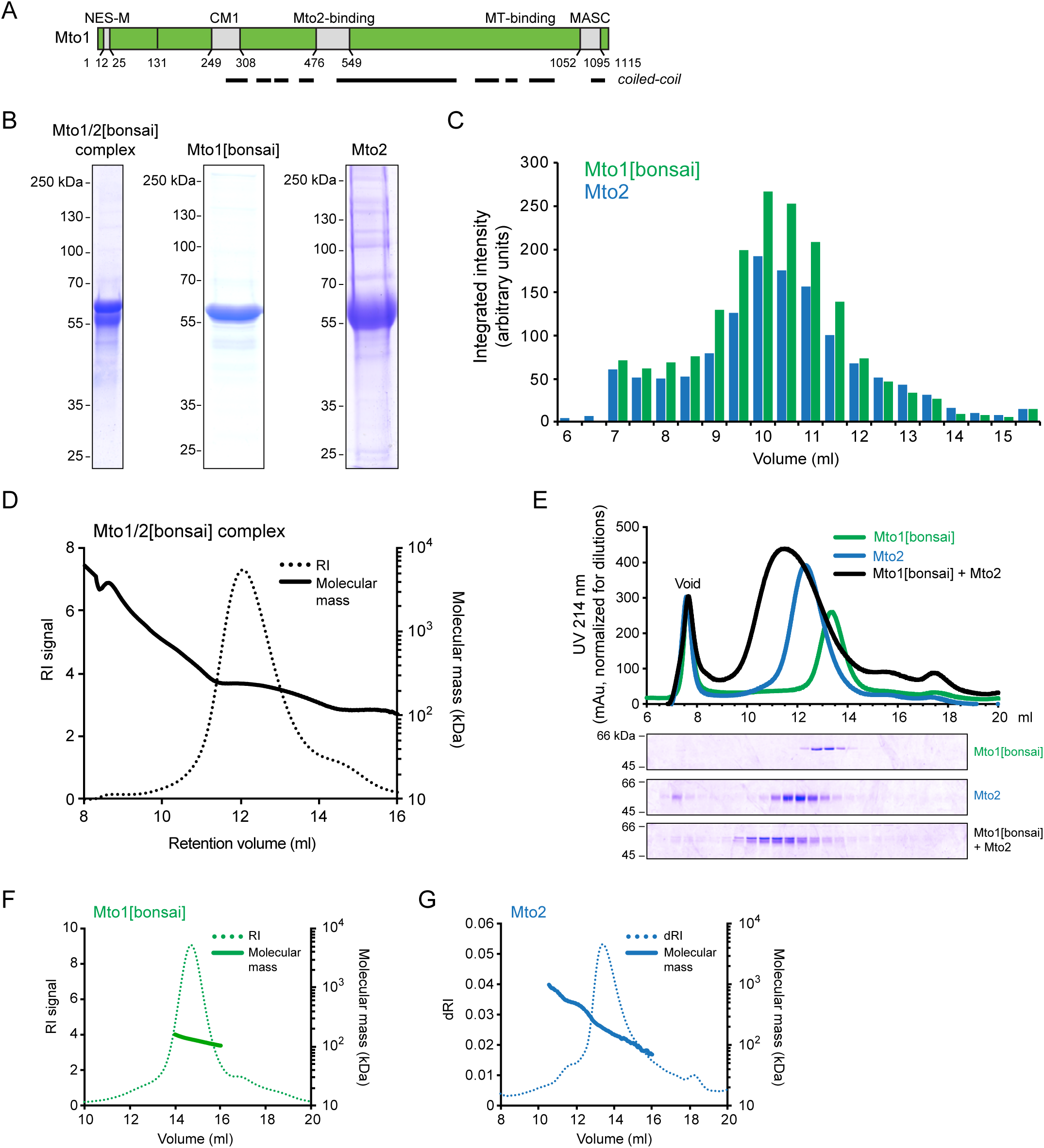
Further characterization of Mto1/2[bonsai] complex, Mto1[bonsai], and Mto2. **(A)** Domain organization of full-length Mto1. Regions are labeled as in Figure 1A. NES-M sequence confers localization to nuclear pore complexes on the nuclear envelope (Bao et al., 2018). MASC region contains modules that target Mto1/2 to the SPB and equatorial MTOC, while boundaries of the microtubule-binding region in the C-terminal region are not completely defined (Samejima et al., 2010). Black lines indicate predicted coiled-coils. **(B)** SDS-PAGE Coomassie Blue (CB) stain of 6xHis-tagged Mto1/2[bonsai] complex, Mto1[bonsai], and Mto2, purified from insect cell expression (see Methods). **(C)** Quantification of SDS-PAGE CB stain of Mto1/2[bonsai] complex after size-exclusion chromatography as shown in Figure 1B. In all fractions, Mto1[bonsai] and Mto2 are present in approximately equal stoichiometry. **(D)** Size-exclusion chromatography with multi-angle laser-light scattering (SEC-MALS) analysis of Mto1/2[bonsai] complex. Sample was injected at ∼4.5 mg/mL. **(E)** Reconstitution of Mto1/2[bonsai] complex by mixing purified Mto1[bonsai] with purified Mto2. SDS-PAGE CB stain is shown underneath. Note the different elution volume and breadth of reconstituted peak. **(F, G)** SEC-MALS analysis of Mto1[bonsai] (F) and Mto2 (G). Samples were injected at ∼1.3 mg/mL (F) and ∼1.6 mg/mL (G). Different Y-axis units (RI signal, dRI) are used for indicating protein peaks in F and G because experiments were performed on different instruments (see Methods). RI indicates refractive index. dRI indicates differential refractive index. Note log_10_ scales for molecular mass in D, F and G.

**Supplementary Figure 2.**
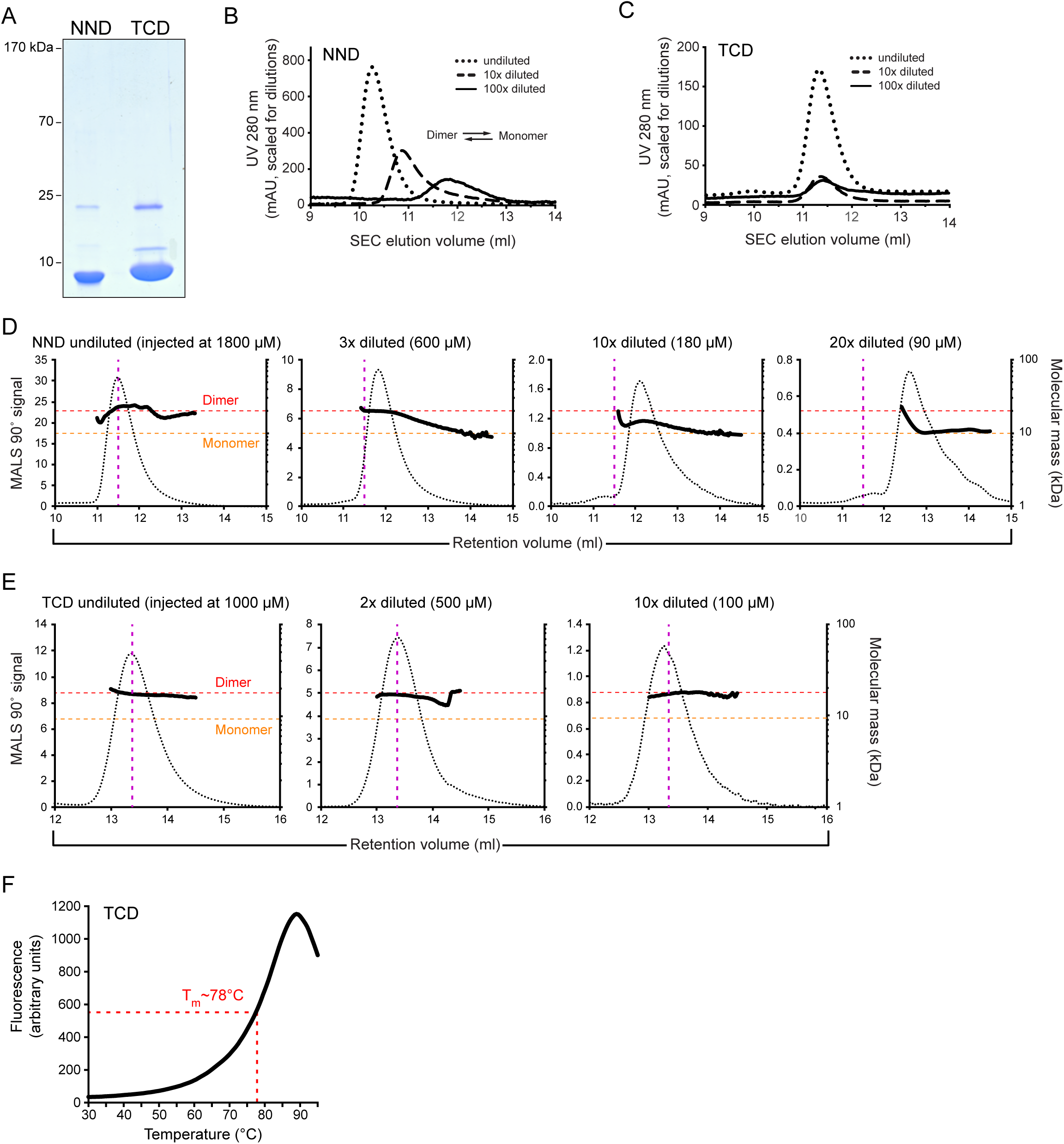
Biophysical characterization of the Mto2 NND and TCD. **(A)** SDS-PAGE Coomassie Blue stain of purified NND and TCD. **(B)** Size-exclusion chromatography (SEC)-dilution analysis of the NND. Upon dilution, the NND shifts to a later-eluting species. **(C)** SEC-dilution analysis of the TCD. Upon dilution, the TCD elution profile is unchanged. **(D)** Size-exclusion chromatography with multi-angle laser-light scattering (SEC-MALS) analysis of the NND, injected into the chromatography column at the different concentrations shown. Red and orange dashed lines indicate dimer and monomer molecular masses, respectively. Purple dashed lines indicate position of elution peak for undiluted NND. Protein elution profiles are shown by MALS 90° signal rather than RI signal because RI is noisy at lower protein concentrations. Data for 600 µM NND are the same as in Figure 2A but plotted using different parameters. **(E)** SEC-MALS analysis of the TCD, injected into the size-exclusion column at different concentrations. Labeling is as in D. Data for 1000 µM TCD are the same as in Figure 2D but plotted using different parameters. **(F)** Thermal denaturation analysis of the TCD. Different peak-elution positions of proteins in different panels (e.g. NND in B vs. D, or TCD in C vs. E) are due to different instruments/methods used. Note log_10_ scales for molecular mass in D and E.

**Supplementary Figure 3.**
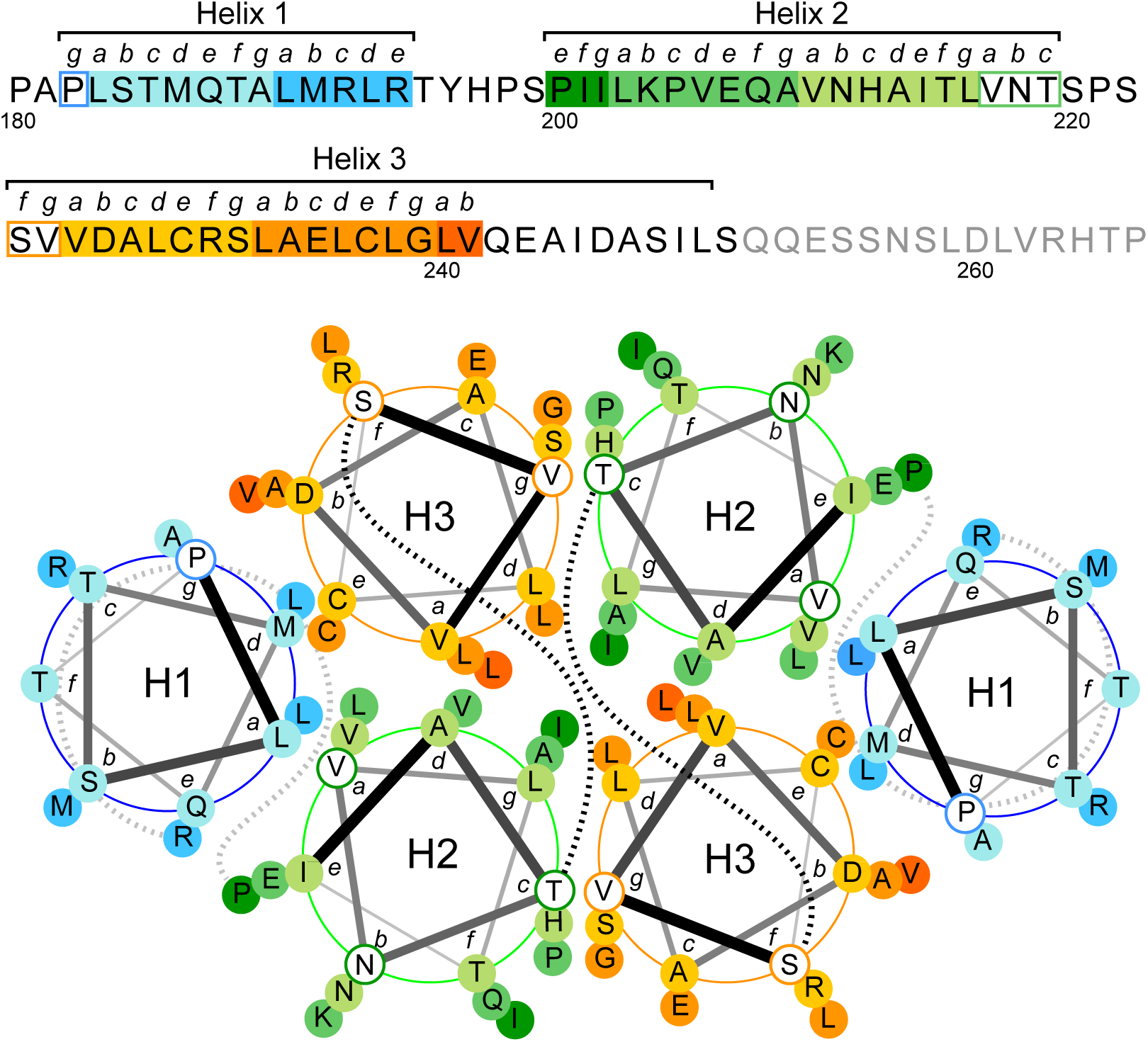
Hydrophobic interactions within the TCD. Helical-wheel plot showing the core of Mto2 TCD dimer (Mto2 residues 182-240), based on X-ray crystal structure in Figure 3. Amino-acid sequence of the entire TCD (residues 180-265) is shown above. Amino-acid residues shown in gray were not observed within the crystal structure in either chain a or chain b. Orientation of the helical-wheel plot is identical to that in the ribbon diagram on the right in Figure 3A. Helix 1 (H1) runs (from N- to C-terminus) into the page. Helix 2 (H2) runs out of the page, towards the viewer. Helix 3 (H3) runs into the page. Within each helix, darker-shaded residues are further away from the viewer. Dashed lines indicate linker residues between helices. Note abundance of hydrophobic residues at interfaces between helices within each monomer, and at the monomer-monomer interface. In particular, aliphatic residues plus methionine are present at “*a*” and “*d*” positions in all helices and at the “*g*” position in Helix 2, and cysteines, which are polar but hydrophobic, are present at the “*e*” position in Helix 3.

**Supplementary Figure 4.**
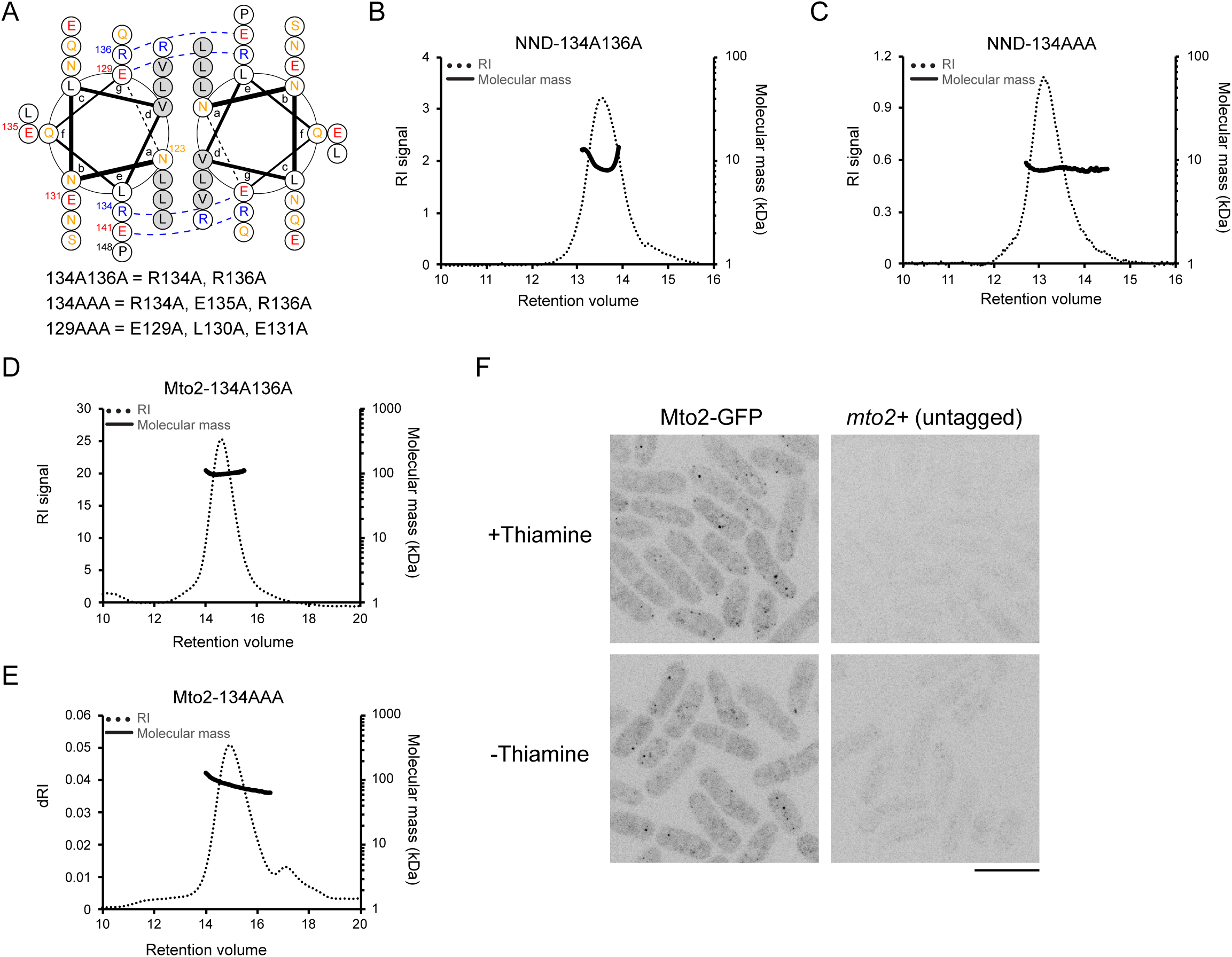
Mutations affecting NND dimerization. **(A)** Helical-wheel plot of the predicted coiled-coil region within the Mto2 NND (residues 123-148). Dashed lines indicate predicted salt bridges. Residues mutated in the 134A136A, 134AAA, and 129AAA mutants are indicated below (the 129AAA mutation is relevant to Figure 6 and Suppl. Figure 6). **(B,C)** SEC-MALS analysis of purified mutant NND proteins NND-134A136A (B) and NND-134AAA (C). Samples were injected at 5.4 mg/mL (B) and 1.3 mg/mL (C). Unlike wild-type NND, which forms homodimers (Figure 2, Suppl. Figure 2), both mutant proteins are monomeric. **(D,E)** SEC-MALS analysis of full-length Mto2 containing the 134A136A (D) and 134AAA (E) mutations. Samples were injected at 1.4 mg/mL (D) and 1.3 mg/mL (E).Unlike wild-type Mto2, which forms higher-order multimers (Figure 1; Suppl. Figure 1), both Mto2-134A136A and Mto2-134AAA are dimeric. Different Y-axis units (RI signal, dRI) are used for indicating protein peaks in D and E because experiments were performed on different instruments (see Methods). **(F)** Additional controls for experiments showing effects of NND and TCD overexpression in *mto1Δ* cells, as in Figure 4C. In *mto1Δ* cells expressing *mto2-GFP* but lacking transgenes, Mto2-GFP puncta are unaffected by the presence vs. the absence of thiamine. In *mto1Δ* cells lacking Mto2-GFP (*mto2+* untagged), neither fluorescent puncta nor cytoplasmic fluorescence is observed. Images were processed identically to those shown in Figure 4C. Scale bar, 10 µm. Note log_10_ scales for molecular mass in B-E.

**Supplementary Figure 5.**
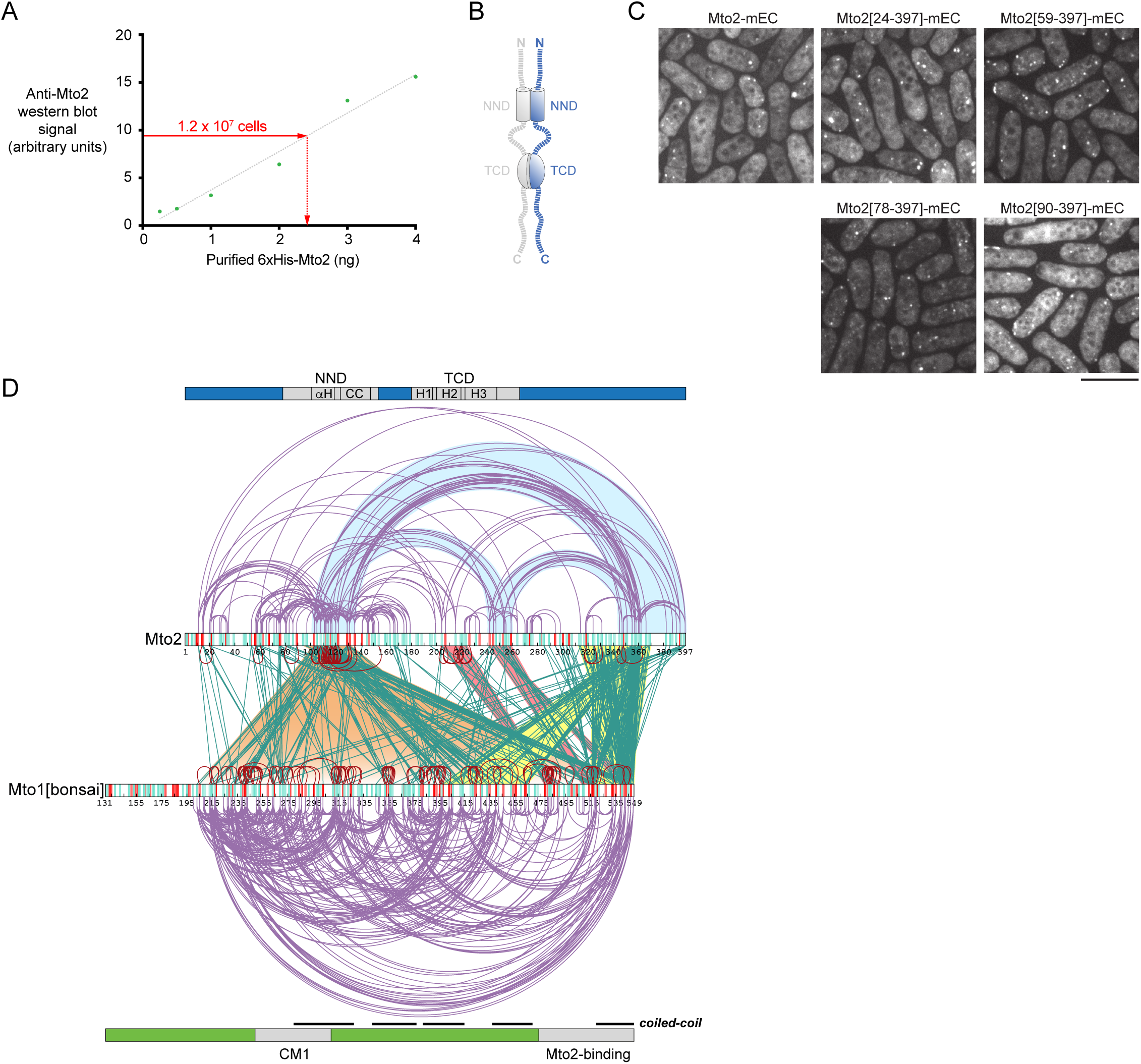
Additional data related to the ABBA multimerization model and CLMS experiments. **(A)** Quantification of endogenous *S. pombe* Mto2 expression. Anti-Mto2 western blot signal from wild-type cell lysate corresponding to 1.2 x 10^7^ cells, compared with western blot signals generated by adding known amounts of purified recombinant Mto2 to *S. pombe mto2Δ* lysate under identical conditions. The derived value of ∼2.0 x 10^-16^ g Mto2 per cell corresponds to a cytosolic concentration of ∼160 nM (see Methods). **(B)** Diagram of a hypothetical “closed bivalent dimer” of Mto2, which might be expected to form if the region between the NND and TCD were completely flexible and the orientation of the NND relative to the TCD were unconstrained. In this scenario, TCD-TCD dimerization would be expected to facilitate NND-NND dimerization by increasing the local concentration of NND. **(C)** *In vivo* puncta formation for wild-type Mto2-mECitrine (Mto2-mEC) and Mto2 N-terminal truncations, in *mto1Δ* strain background. Puncta are seen with all truncations. Scale bar, 10 µm. **(D)** Complete set of Mto1-Mto2, Mto2-Mto2, and Mto1-Mto1 interactions identified by EDC crosslinking of Mto1/2[bonsai] complex. Data are copied from Figure 5C but also include the Mto1-Mto1 crosslinks that could be either inter- or intramolecular; for clarity these were not shown in Figure 5C.

**Supplementary Figure 6.**
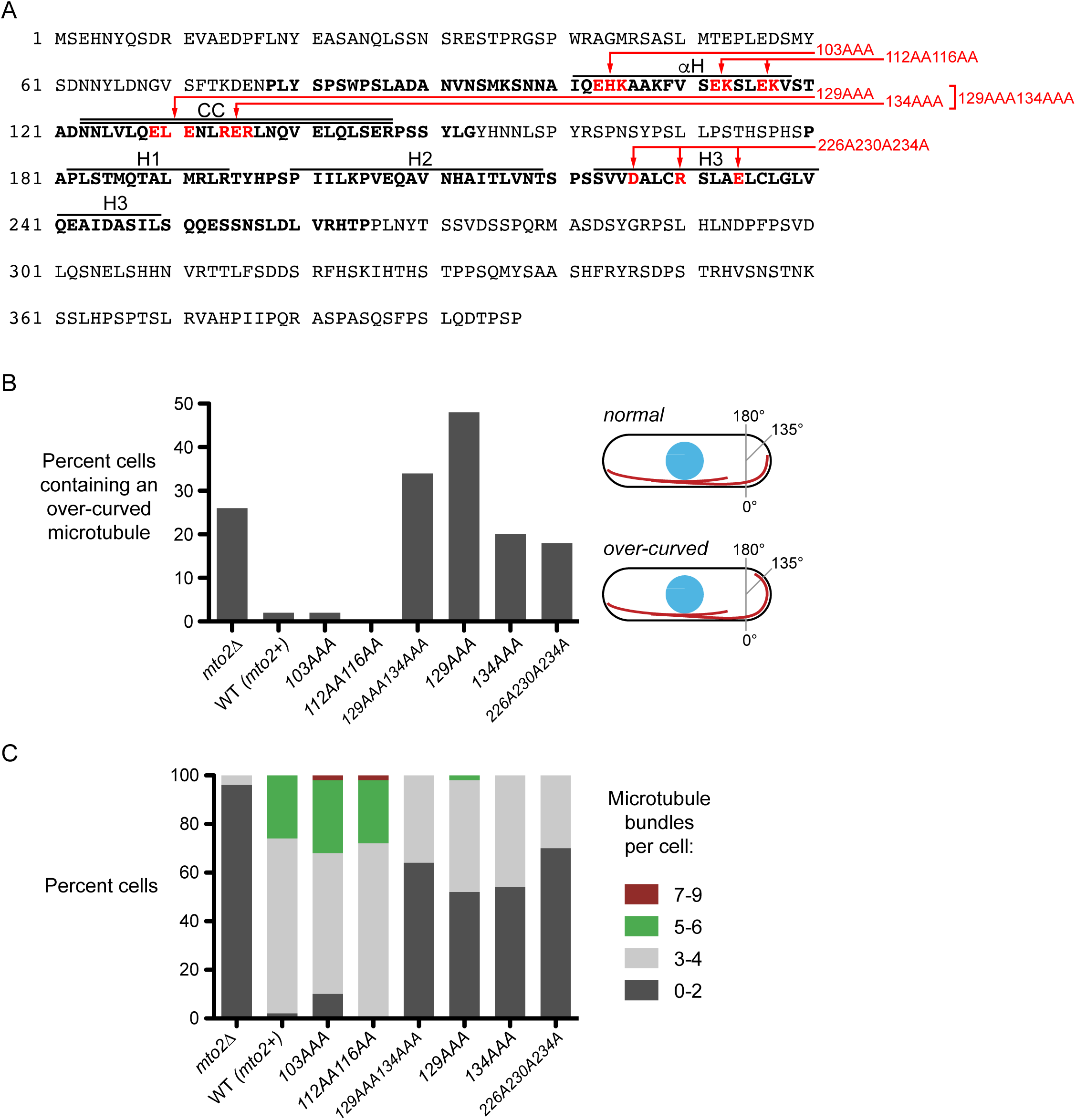
Quantification of microtubule phenotypes in Mto2 NND and TCD mutants. **(A)** Mto2 sequence, showing position of mutations described in Figure 6. Bold text indicates residues in NND (78-153) and TCD (180-265). Helical regions are labeled as in Figure 2. **(B)** Quantification of over-curved microtubules in wild-type (WT, *mto2*+) cells and in the *mto2* mutants indicated, from images as in Figure 6A. An over-curved microtubule is defined as having a bending angle at the cell tip greater than 135° (see diagram). For each strain, n=50 cells scored. **(C)** Number of microtubule bundles per cell in wild-type cells and in the *mto2* mutants indicated, from images as in Figure 6A. For each strain, n=50 cells scored.

**Supplementary Figure 7.**
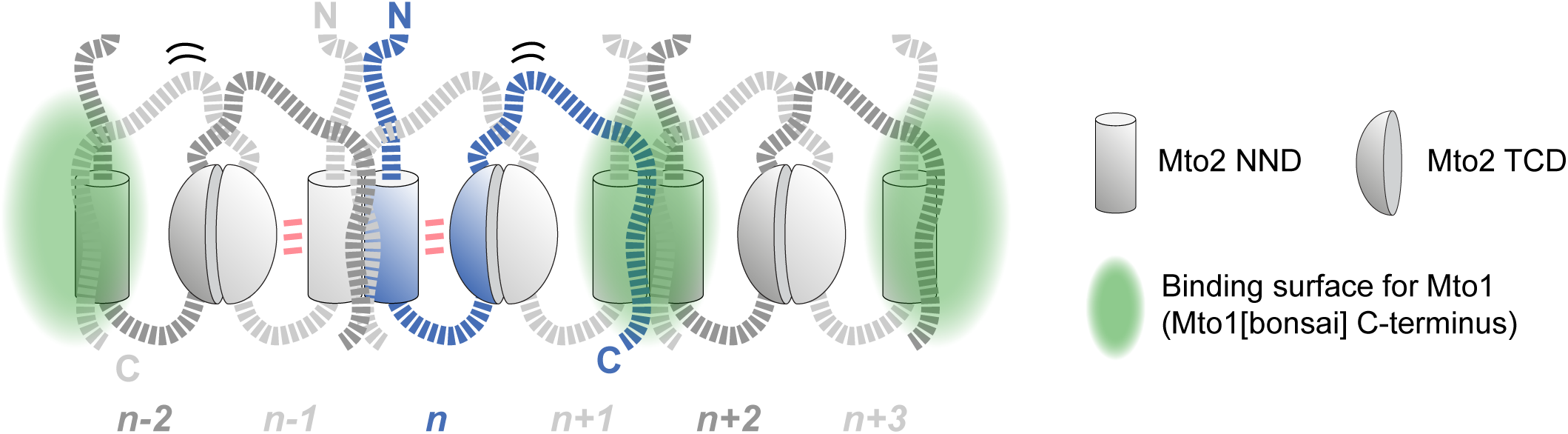
Alternative model for Mto1/2 complex architecture. In this variant of the model in Figure 7, Mto2 C-terminus interacts with a different NND-NND interface from that shown in Figure 7 (e.g. the C-terminus of Mto2*_n_* interacts with the NND-NND interface between Mto2*_n+1_* and Mto2*_n+2_*, rather than between Mto2*_n_* and Mto2*_n-1_*. For this model to be plausible, the constraints on NND and TCD orientation that prevent closed bivalent dimer formation (Suppl. Figure 5) would likely be conferred by the linker sequence between the NND and the TCD rather than by the C-terminus interacting with the NND (see Figure 7). Although this linker is predicted to be intrinsically disordered, it might adopt a specific structure in the presence of the NND and/or TCD.

